# Bacterial Peptidoglycan Extends Lifespan by Activating Lysosomal Activity through V-ATPase Binding

**DOI:** 10.64898/2026.08.28.747948

**Authors:** Herui Fu, Jia Tie, Yan Zhou, Xin Wang, Zhao Shan, Bin Qi

## Abstract

Lysosomal dysfunction is a hallmark of aging, yet whether microbial components actively regulate this organelle to influence longevity remains unknown. Here, we identify bacterial peptidoglycan (PGN), a major cell wall component degraded by host lysozyme, as an evolutionarily conserved activator of lysosomal function that extends lifespan in both *C. elegans* and mice. We show that aging leads to an intestinal decline in lysozyme expression, which impairs bacterial cell-wall digestion and results in systemic PGN deficiency. Late-life PGN supplementation (starting at 18 months of age) significantly prolongs mouse lifespan and improves healthspan. Mechanistically, PGN localizes to lysosomes and directly binds V-ATPase subunits, enhancing ATP hydrolysis activity and promoting lysosomal acidification. This effect is abolished by V-ATPase inhibition (bafilomycin A1) or genetic disruption of lysosomal components (*cup-5* and *vha-12* mutants), confirming that functional V-ATPase is strictly required for lysosomal function and the longevity benefit. Importantly, PGN restores lysosomal acidification in aged cells, alleviates cellular senescence markers, and improves multiple hallmarks of aging including locomotion and muscle integrity. Collectively, these findings reveal an evolutionarily conserved mechanism whereby hosts exploit bacterial cell wall components to maintain cellular homeostasis, establishing a gut microbiome–lysosome–longevity axis with implications for microbiome-based anti-aging interventions.

**Highlights:**

- Aging reduces intestinal lysozyme activity, depleting systemic PGN bioavailability
- PGN binds V-ATPase subunits, enhancing ATP hydrolysis and lysosomal acidification
- Bacterial PGN supplementation extends lifespan and improves healthspan
- PGN maintains multi-tissue lysosomal homeostasis and reduces cellular senescence during aging

## Introduction

Aging is a complex biological process characterized by the progressive deterioration of cellular and physiological integrity, leading to heightened susceptibility to disease (López-Otin et al., 2023; López-Otín et al., 2013). Among the organelles affected, lysosomes—the cell’s primary catabolic and recycling hubs—are critically implicated in this process (Ballabio and Bonifacino, 2020; Folick et al., 2015; Ramachandran et al., 2019; Steinhauser et al., 2026; Sun et al., 2020). Accumulating evidence positions lysosomal dysfunction as a pivotal driver in the pathogenesis of various age-related disorders, including Alzheimer’s disease, Parkinson’s disease, and sarcopenia (Carmona-Gutierrez et al., 2016; Lee et al., 2010). Consequently, the therapeutic restoration of lysosomal homeostasis has emerged as a promising anti-aging strategy (Platt et al., 2018; Steinhauser et al., 2026). Functionally, lysosomal activity relies on an acidic luminal pH (4.5–5.0), a gradient stringently maintained by the vacuolar-type H^+^-ATPase (V-ATPase), which couples ATP hydrolysis to proton translocation across the membrane (Forgac, 2008). Notably, V-ATPase activity declines with advancing age, a loss that exacerbates lysosomal failure and contributes to neurodegenerative pathology (Colacurcio and Nixon, 2016). Despite its established pathological importance, no naturally occurring, exogenously administrable molecule capable of directly activating V-ATPase and rejuvenating lysosomal function has been identified to date.

The gut microbiome has emerged as a pivotal modulator of host aging trajectories (Bana and Cabreiro, 2019). Gut dysbiosis is formally designated as one of the twelve hallmarks of aging (López-Otín et al., 2013), with age-related compositional shifts known to drive systemic inflammation and metabolic dysfunction (Ghosh et al., 2020; Thevaranjan et al., 2017). Intriguingly, a study in a Chinese population revealed that the intestinal microbial community of centenarians shares compositional features with that of younger individuals (Pang et al., 2023). In the turquoise killifish, aging correlates with a marked loss of microbial diversity, and transplantation of gut microbes from young donors into aged conspecifics effectively attenuates age-related decline (Smith et al., 2017). Likewise, in *Caenorhabditis elegans*, specific bacterial isolates, mutant strains, or their metabolic products have been demonstrated to extend lifespan (Gusarov et al., 2013; Han et al., 2017). Yet the evolutionarily conserved molecular signals through which the microbiota communicates with host cells to modulate aging remain incompletely characterized, and the mechanisms by which individual microbial-derived molecules maintain intracellular lysosomes homeostasis are essentially unknown.

Peptidoglycan (PGN) constitutes the primary structural scaffold of the bacterial cell wall, comprising glycan chains of alternating N-acetylglucosamine (NAG) and N-acetylmuramic acid (NAM) cross-linked by short peptide stems (Vollmer et al., 2008). Within the intestinal lumen, PGN is liberated from resident bacteria and subsequently degraded by lysozymes—the principal innate immune effectors secreted by Paneth cells (Bevins and Salzman, 2011). PGN fragments, most notably muramyl dipeptide (MDP), are recognized by the cytosolic pattern recognition receptors NOD1 and NOD2, thereby activating NF-κB-dependent inflammatory signaling (Girardin et al., 2003). Previous studies by Schwarzer et al. (2023) demonstrated that microbe-mediated intestinal NOD2 stimulation promotes linear growth in undernourished infant mice (Schwarzer et al., 2023), and Gabanyi et al. (2022) showed that neuronal Nod2-dependent bacterial sensing regulates appetite and body temperature (Gabanyi et al., 2022), both studies highlight PGN’s physiological roles through classical immune receptor (NOD2) signaling. More recently, PGN muropeptides have been demonstrated to promote mitochondrial homeostasis by functioning as agonists of ATP synthase in *C. elegans* intestinal cells—an effect that has since been validated in mammals (Tian et al., 2024; Tian and Han, 2022). Additionally, PGN has been reported to induce protective mitophagy by binding p62 in the mice liver (Tie et al., 2025) and to stimulate food digestion via inhibition of the unfolded protein response associated with mitochondria (UPRmt) in *C. elegans* (Hao et al., 2024). However, whether PGN exerts functions beyond immune activation and mitochondrial regulation—particularly in modulating host aging through lysosomal pathways—remains entirely unexplored.

Here we report that peptidoglycan (PGN) functions as an evolutionarily conserved microbiota-derived longevity signal that directly engages V-ATPase V1 subunits to enhance lysosomal acidification and extend healthy lifespan in both *C. elegans* and mice. More broadly, our study introduces a conceptual framework in which a bacterial cell-wall polymer functions as a physiological regulator of a core eukaryotic organelle. By identifying lysosomal V-ATPase as a direct target of a microbiota-derived molecule, our work provides a molecular mechanism linking microbial cell-wall metabolism to host lysosomal homeostasis and healthy aging. We anticipate that this framework will stimulate new investigations into host–microbiota communication and inspire therapeutic strategies aimed at restoring lysosomal function in aging and age-associated diseases.

## Results

### Aging impairs intestinal bacterial digestion and reduces PGN bioavailability

To investigate whether aging affects intestinal bacterial processing, we monitored the accumulation of GFP-labeled *E. coli* (OP50-GFP) in *C. elegans* at different ages. GFP fluorescence intensity in the intestinal lumen increased significantly during aging (Figure 1A), indicating a progressive decline in the capacity of aging worms to digest ingested bacteria.

**Figure 1.**
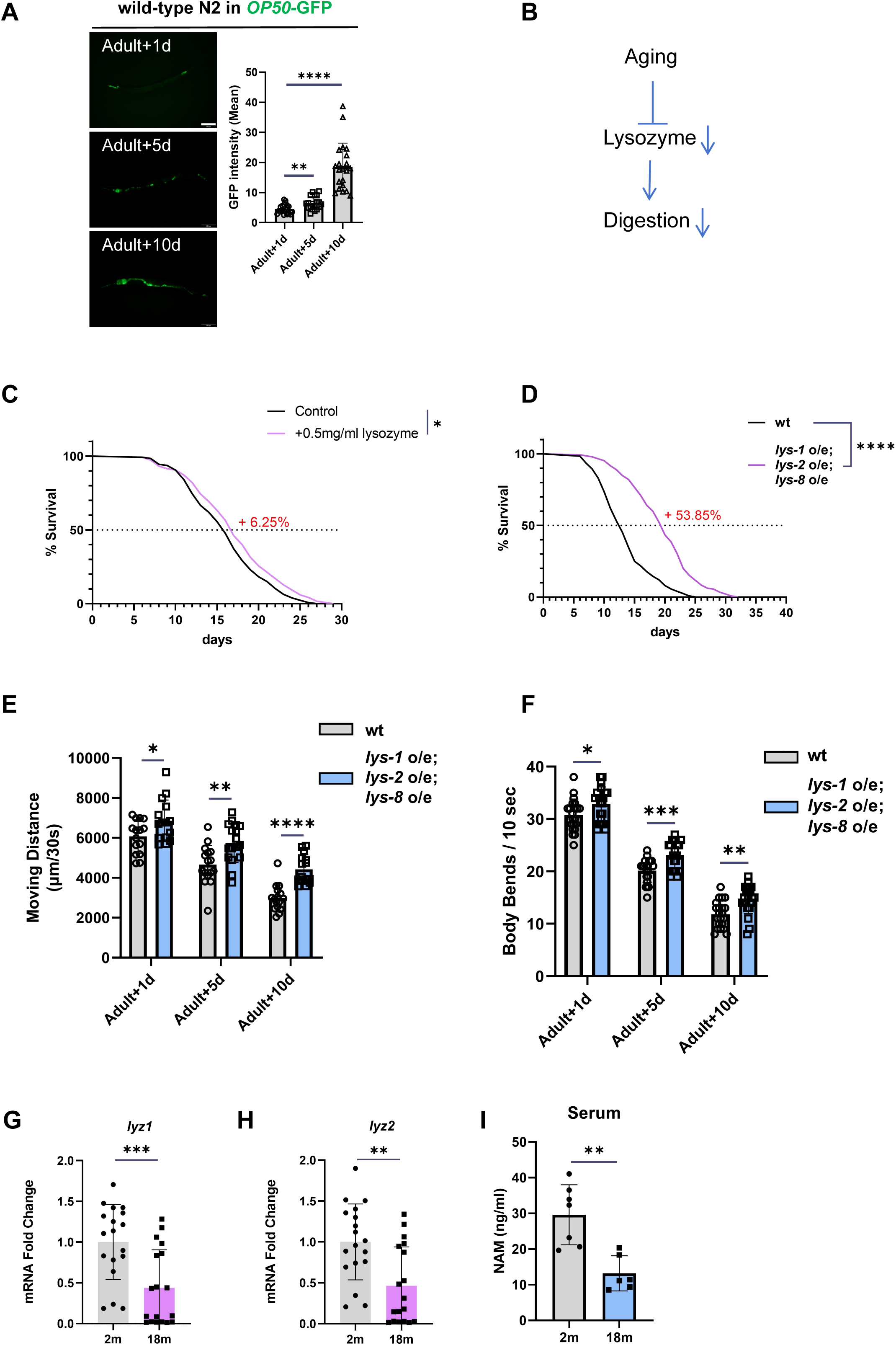
Aging impairs intestinal bacterial digestion and lysozyme activity, and lysozyme overexpression extends lifespan. (A) Fluorescence images and quantification of OP50-GFP bacteria in the *C. elegans* intestinal lumen at Adult+1d, Adult+5d, and Adult+10d. Scale bar, 200 μm. Data are mean ± SD; Statistical analysis were performed by one-way ANOVA (**P < 0.01, ****P < 0.0001). (B) Schematic depicting the relationship between aging, declining lysozyme activity, and impaired bacterial digestion. (C) Survival curves of wild-type *C. elegans* fed OP50 with or without 0.5 mg/ml exogenous lysozyme. Lysozyme supplementation extends median lifespan by +6.25% [*P < 0.05, by log-rank (Mantel-Cox) test; n ≥ 100 worms per replicate; N=3 independent biological replicates] . (D) Survival curves of wild-type (wt) and *lys-1 o/e; lys-2 o/e; lys-8 o/e* triple overexpression transgenic worms. Lysozyme overexpression extends median lifespan by +53.85% [****P < 0.0001, by log-rank (Mantel-Cox) test; n ≥ 100 worms per replicate; N=3 independent biological replicates]. **(E, F)** Moving distance (E) and body bends (F) of wild-type (wt) and triple lysozyme overexpression (*lys-1, lys-2, lys-8* OE) animals at Adult+1d, Adult+5d, and Adult+10d. *P < 0.05, **P < 0.01, ***P < 0.001, ****P < 0.0001, by unpaired two-tailed Student’s t-tests. n ≥ 15 worms per replicate; N= 3 independent biological replicates. **(G, H)** Bar graphs showing qPCR-quantified mRNA levels of *lyz1* (G) and *lyz2* (H) in ileal tissue from 2-month-old and 18-month-old mice (n ≥ 5 mice per group; 3 technical replicates each). Data are mean ± SD. ***P < 0.001, **P < 0.01(by unpaired two-tailed Student’s t-tests). **(I)** Bar graphs showing serum NAM levels measured by chromatography as a proxy for systemic PGN bioavailability in 2-month-old versus 18-month-old mice (n ≥ 5 mice per group). Data are mean ± SD. **P < 0.01; ns, not significant (by unpaired two-tailed Student’s t-tests). See also Figure S1, Figure S2 and Table S1.

Lysozymes are antimicrobial enzymes that cleave bacterial cell walls, releasing peptidoglycan (PGN) as a degradation product. We hypothesized that reduced lysozyme expression might underlie this age-related decline in bacterial digestion (Figure 1B). Published data (Gao et al., 2024; Kong et al., 2024) indicate that *lys-1*, *lys-2*, and *lys-8* show the most pronounced reduction among lysozyme-related genes in aged worms (day 12) (Figure S1A-S1B). To test whether enhancing lysozyme activity could extend lifespan, we first supplemented worm culture medium with exogenous lysozyme at 0.5 mg/ml, which significantly extended median lifespan by 6.25% (Figure 1C). Furthermore, we generated transgenic worms simultaneously overexpressing *lys-1, lys-2,* and *lys-8*; these animals showed a 53.85% increase in median lifespan compared to wild-type controls (Figure 1D) and exhibited significantly greater locomotor activity (distance moved within 30 seconds, Figure 1E) and higher body bending frequency (Figure 1F) throughout life, indicating improved healthspan. Bacterial accumulation in the gut was also reduced in these transgenic animals from Adult+1d to Adult+10d (Figure S1C). These results demonstrate that aging-associated decline in lysozyme expression contributes to reduced digestive capacity and that restoration of lysozyme levels can extend both lifespan and healthspan.

Paneth cells are the principal source of C-type lysozyme, a β-1,4-N-acetylmuramoylhydrolase that enzymatically processes bacterial cell wall (Yu et al., 2020). Published data demonstrate that lysozyme, a marker of Paneth cells, is strongly downregulated in the ileum of aged (19-month-old) mice compared with young (10-week-old) controls (Sovran et al., 2019). Similarly, in our own analysis, aging in mammals was associated with reduced lysozyme expression: mRNA levels of both *lyz1* and *lyz2* were significantly lower in ileal tissue of 18-month-old mice relative to 2-month-old animals (Figures 1G, 1H). Consistent with diminished lysozyme-mediated PGN degradation, serum concentrations of N-acetylmuramic acid (NAM)—a diagnostic marker of PGN breakdown—were markedly decreased in aged mice (Figure 1I, Figure S2A-C).

Together, these data indicate that the age-dependent decline in intestinal lysozyme is evolutionarily conserved and results in reduced systemic availability of PGN-derived metabolites. These findings establish intestinal lysozyme activity as a limiting factor for longevity and suggest that PGN products generated from bacterial cell wall hydrolysis may serve as pro-longevity signals.

### PGN supplementation extends lifespan and improves healthspan in *C. elegans*

Given that lysozymes cleave bacterial cell walls to release peptidoglycan (PGN) (Figure 2A, 2B), and that aging reduces lysozyme-mediated PGN production (Figure 1I, 2B), we hypothesized that exogenous PGN supplementation might recapitulate the pro-longevity effects of enhanced bacterial digestion (Figure 2B). To test this, we supplemented the standard *E. coli* (OP50) diet with purified PGN derived from *Bacillus subtilis* (B.s-PGN) (Figure S3A). PGN supplementation extended median lifespan by approximately 10% compared to OP50-fed controls (Figure 2C). In locomotion assays, PGN-treated worms exhibited significantly greater moving distances (Figure 2D, Figure S3B) and higher body bending frequencies (Figure 2E) at the Adult+10d time point, when age-related decline in wild-type animals is most pronounced, indicating that PGN treatment improved multiple healthspan parameters.

**Figure 2.**
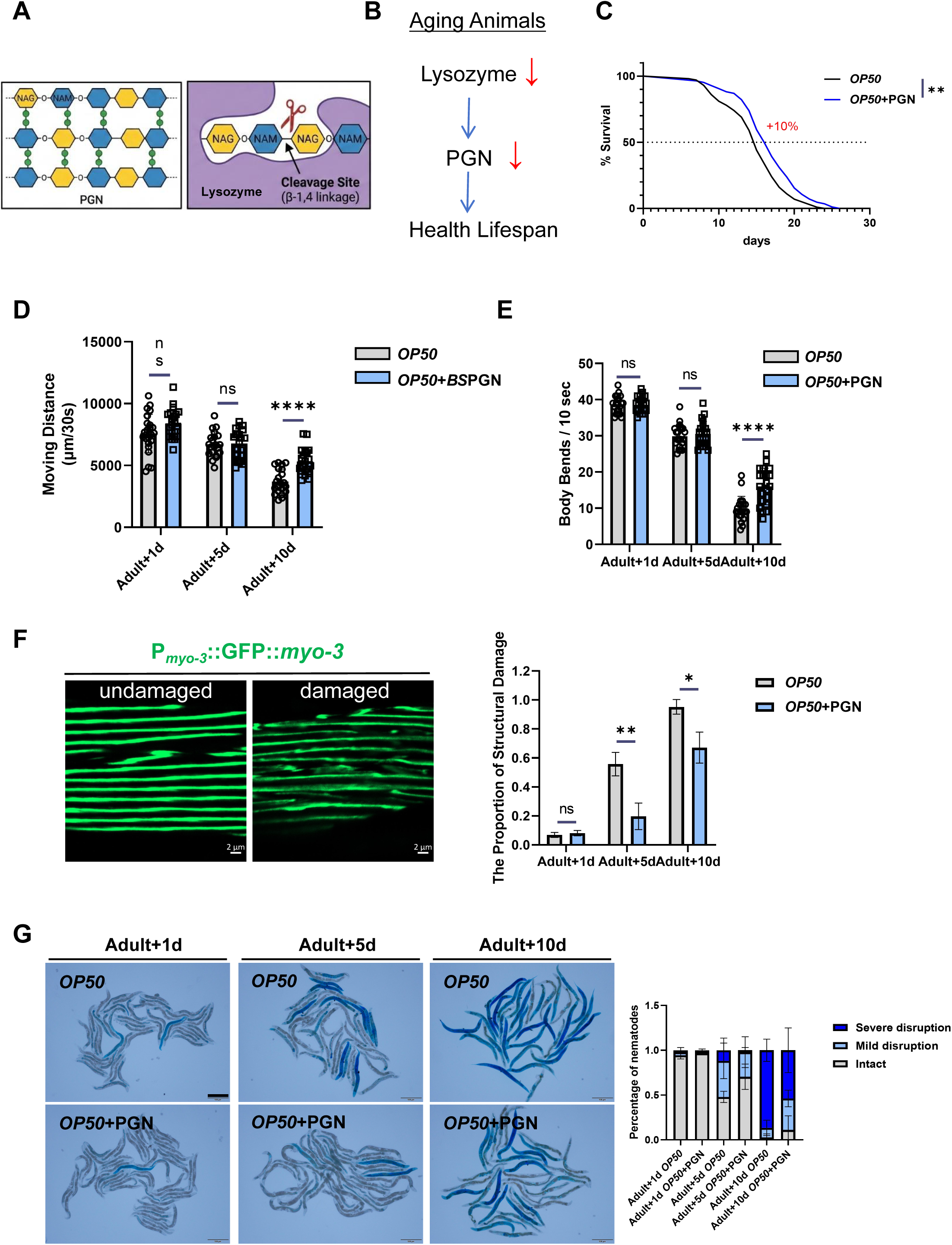
PGN supplementation extends lifespan and improves age-related functional decline in *C. elegans*. **(A)** Schematic of PGN chemical structure showing NAG-NAM disaccharide repeating units and the lysozyme cleavage site at the β-1,4 glycosidic bond. **(B)** Schematic of the proposed pathway: aging reduces lysozyme activity, leading to reduced PGN release and impaired healthspan. **(C)** Survival curves of *C. elegans* fed OP50 or OP50 supplemented with B.s-PGN. PGN extends median lifespan by +10% (**P < 0.01 by log-rank (Mantel-Cox) test ; n ≥ 100 worms per replicate; N=3 independent biological replicates). **(D, E)** Moving distance (D) and body bends (E) at Adult+1d, Adult+5d, and Adult+10d for OP50 and OP50+B.s-PGN animals. Data are mean ± SD. ****P < 0.0001; ns, not significant (by unpaired two-tailed Student’s t-tests). n ≥ 15 worms per replicate; N= 3 independent biological replicates. **(F)** Representative fluorescence images of P*myo-3*::GFP::myo-3 reporter in undamaged and damaged muscle categories. Bar graph quantifies the proportion of worms with structural muscle damage at Adult+1d, Adult+5d, and Adult+10d. Data are mean ± SD. ****P < 0.0001; ns, not significant (by unpaired two-tailed Student’s t-tests). (n ≥ 20 worms per replicate; N = 3 replicates). Scale bar, 2 μm. **(G)** Representative images and quantification of intestinal barrier integrity (Smurf assay) showing proportions of worms classified as intact, mildly disrupted, or severely disrupted at Adult+1d, Adult+5d, and Adult+10d in OP50 and OP50+B.s-PGN conditions. Data are mean ± SD. n ≥ 20 worms per replicate; N= 3 independent biological replicates. See also Figure S3 and Table S1.

Aging is accompanied by progressive deterioration of muscle structure. We assessed muscle integrity using the P*myo-3*::GFP::myo-3 reporter, which labels body wall muscle fibers. At both Adult+5d and Adult+10d, PGN-treated animals showed a significantly lower proportion of worms with structurally damaged muscle fibers compared to controls (Figure 2F, Figure S3C), indicating that PGN treatment effectively delayed age-related muscle structural decline. Furthermore, intestinal barrier integrity, a hallmark of gut health that declines with age, was assessed by the Smurf assay (Rera et al., 2012). PGN supplementation maintained intestinal permeability barrier function during aging (Figure 2G).

Collectively, these results demonstrate that bacterial-derived PGN is a potent longevity-promoting factor that extends both lifespan and healthspan in *C. elegans*.

### PGN dietary supplementation extends lifespan and promotes healthy aging in mice

To determine whether the pro-longevity effects of PGN are conserved in mammals, we initiated PGN dietary supplementation in 18-month-old aged mice — a late-in-life stage corresponding to advanced age in humans (Figure 3A). PGN treatment caused a significant lifespan extension in both sexes. In male mice, median lifespan was extended by 9.74% (control: 847 days, PGN: 929.5 days; P=0.0047, log-rank test) and maximal lifespan [defined as the age at 90% mortality in the joint lifespan distribution, according to Wang et al., 2004(Wang et al., 2004)] was extended by 21%. In female mice, median lifespan was extended by 11% (control: 892 days, PGN: 990.5 days; P=0.0083, log-rank test) and maximal lifespan was extended by 12.7%. Importantly, body weight and food intake in PGN-treated groups were not decreased compared to control groups (Figure S4A–S4D), which rules out the possibility that the observed lifespan extension is mediated by caloric restriction (Schmauck-Medina et al., 2026).

**Figure 3.**
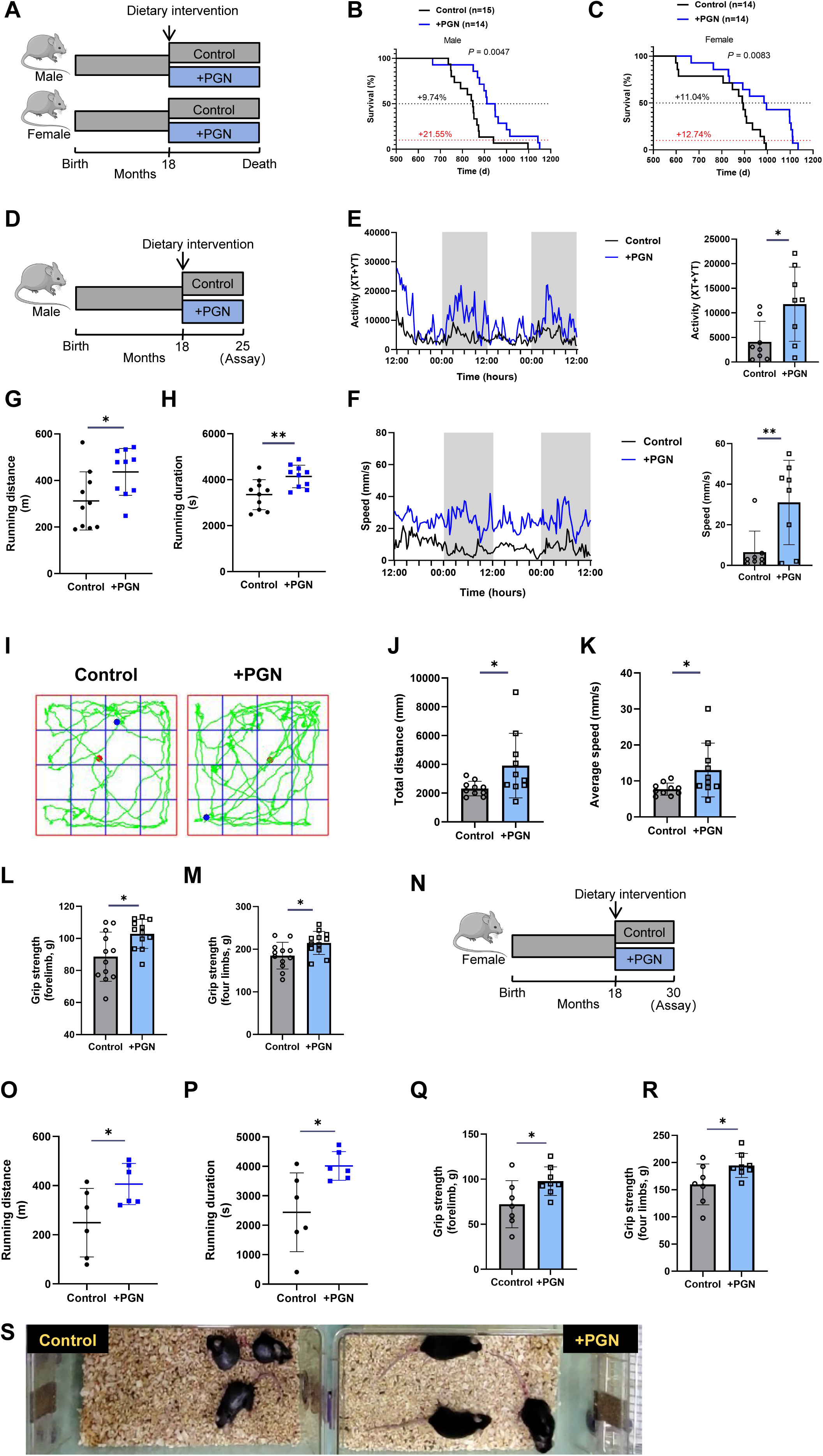
PGN dietary supplementation extends lifespan and improves healthspan in aged mice. **(A)** Dietary intervention schematic: PGN or control chow was initiated at 18 months of age in both male and female cohorts and maintained until natural death. **(B, C)** Survival curves of male (B; n = 14–15 per group; P = 0.0047) and female (C; n = 14 per group; P = 0.0083) mice on control or PGN-supplemented diet. Log-rank (Mantel-Cox) test. **(D)** Schematic of the healthspan assessment protocol: PGN dietary intervention was initiated at 18 months in male mice; behavioral and functional tests were performed at 25 months. **(E, F)** Metabolic cage analysis showing 48 h locomotion activity (E) and locomotion speed (F) over the monitoring cycle (shaded regions, dark phase). Bar graphs show mean values during the midpoint of the dark phase (n = 8 mice per group). Data are mean ± SD. *P < 0.05, **P < 0.01 (by unpaired two-tailed Student’s t-tests). **(G, H)** Treadmill running distance (G) and running duration (H) in 25-month-old male mice (n = 10 per group). Data are mean ± SD. *P < 0.05, **P < 0.01 (by unpaired two-tailed Student’s t-tests). **(I)** Representative open-field traces from control and +PGN mice. **(J, K)** Open-field total distance traveled (J) and average speed (K) (n = 10 mice per group). Data are mean ± SD. *P < 0.05 (by unpaired two-tailed Student’s t-tests). **(L, M)** Forelimb (L) and four-limb (M) grip strength (n ≥ 10 per group; each data indicates average taken from 3 measurements/mouse). Data are mean ± SD.*P < 0.05 (by unpaired two-tailed Student’s t-tests). **(N)** Schematic diagram of the experimental design for PGN dietary intervention in female mice. Female mice were fed a PGN-supplemented diet starting at 18 months of age and maintained until 30 months of age, at which point various physiological parameters were uniformly assessed. **(O, P)** Treadmill running distance (O) and running duration (P) in 30-month-old female mice (n = 6 per group). Data are mean ± SD. *P < 0.05 (by unpaired two-tailed Student’s t-tests). **(Q, R)** Forelimb (Q) and four-limb (R) grip strength (n ≥ 6 mice per group; each data indicates average taken from 3 measurements/mouse). Data are mean ± SD.*P < 0.05 (by unpaired two-tailed Student’s t-tests). **(S)** Representative images of female mice treated with PGN. See also Figure S4 and Table S2.

To assess the effects of PGN on healthspan, we performed comprehensive functional evaluations in male mice after 7 months of PGN treatment, at 25 months of age (Figure 3D). Metabolic cage analysis revealed that PGN-treated mice exhibited significantly higher spontaneous activity levels and locomotion speed throughout the 48-hour monitoring cycle (Figures 3E, 3F). Treadmill testing demonstrated that PGN-treated mice ran significantly longer distances (Figure 3G) and for longer durations (Figure 3H). Open field testing confirmed that PGN-supplemented mice traveled significantly greater total distances (Figures 3I, 3J) and at higher average speeds (Figure 3K) compared to controls. Additionally, grip strength measured for both forelimb and all four limbs was significantly enhanced in PGN-treated animals (Figures 3L, 3M).

We further validated these findings in female mice. Female mice were treated with PGN from 18 months of age and assessed at 30 months of age following 12 months of treatment (Figure 3N). Treadmill testing showed that PGN-treated females ran significantly longer distances (Figure 3O) and durations (Figure 3P), and grip strength for both forelimbs and all four limbs was significantly increased (Figure 3Q-3R). Moreover, compared to controls, PGN-treated females at 28 months of age exhibited significantly higher spontaneous activity levels (Video S1), a more lustrous and uniform fur coat, and reduced alopecia (Figure 3S).

Collectively, these data establish that PGN supplementation extends both lifespan and healthspan in aged mice, ameliorating multiple hallmarks of physical frailty.

### PGN localizes to lysosomes

Given that lysosomal acidity and degradation activity decline during aging in *C. elegans* (Sun et al., 2020), we asked whether this deterioration is conserved in mammals. To test this, we measured lysosomal acidity in primary hepatocytes using LysoSensor staining and found that fluorescence intensity in cells from 18-month-old mice was significantly lower than that in cells from 2-month-old mice (Figure 4A), directly confirming that aging reduces hepatocyte lysosomal acidification capacity and suggesting a progressive loss of lysosomal function with age.

**Figure 4.**
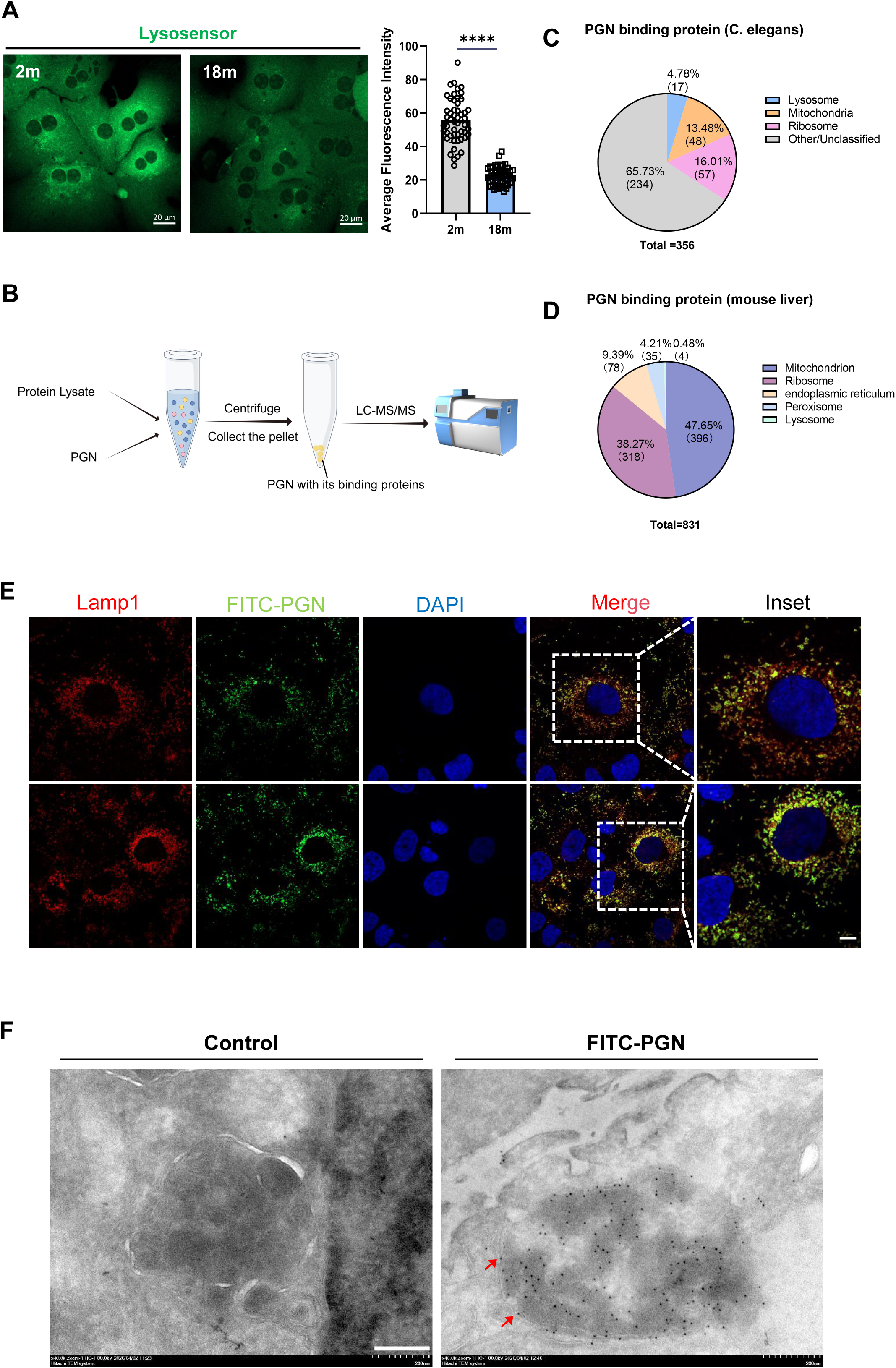
PGN binding partners are enriched for lysosomal proteins, and PGN localizes to lysosomes in mammalian cells. **(A)** LysoSensor Green fluorescence images and quantification in primary hepatocytes from 2-month-old and 18-month-old mice, demonstrating age-dependent decline in lysosomal acidity. Scale bar, 20 μm. Data are mean ± SD. ****P < 0.0001 (by unpaired two-tailed Student’s t-tests). n ≥ 20 cells per replicate; N= 3 independent biological replicates. **(B)** Schematic of the protocol for assessing PGN-binding proteins. **(C, D)** Pie charts showing functional categories of PGN-binding proteins identified by co-precipitation mass spectrometry from *C. elegans* (A; total = 356 proteins) and mouse liver (B; total = 831 proteins). **(E)** Representative confocal immunofluorescence images of Huh7 cells co-incubated with FITC-PGN (green) and immunostained for LAMP1 (red, lysosomal marker), with DAPI (blue) for nuclei. Merged image and inset show extensive co-localization (yellow). Scale bar, 20 μm. **(F)** Transmission electron microscopy images of control and FITC-PGN-treated Huh7 cells with immunogold labeling (anti-FITC, 6 nm gold particles). Gold signals (Arrow) are specifically detected within lysosomal structures in FITC-PGN-treated cells. Scale bar, 200 nm. See also Figure S5 and Table S3-S5.

Given that PGN supplementation extends both lifespan and healthspan in worms and mice, we next investigated whether lysosomes serve as the functional target through which PGN exerts its longevity-regulating effects. To this end, we first performed screening for PGN-binding proteins using co-precipitation coupled with mass spectrometry: total protein lysates from *C. elegans* and mouse liver were incubated with bacterial-derived PGN in vitro, and interacting proteins were identified (Figure 4B). Functional enrichment analysis revealed a subset of proteins related to lysosomal activity regulation in both species (Figure 4C-4D; Table S3-S4). Moreover, published data (Tian and Han, 2022) have also shown that PGN binds to several *C. elegans* lysosomal proteins (Figure S5; Table S5). Although this subset constituted a relatively modest proportion of the total interactome, the data suggested that PGN may target lysosomes by binding to specific lysosome-associated proteins.

To verify the direct spatial association between PGN and lysosomes, we first examined whether PGN could be internalized by cells and trafficked to lysosomes. FITC-labeled PGN was co-incubated with Huh7 hepatic cells, and confocal immunofluorescence microscopy revealed extensive co-localization of FITC-PGN with the lysosomal marker LAMP1 (Figure 4E), indicating that internalized PGN is directed to the lysosomal compartment. To corroborate this at the ultrastructural level, we performed transmission electron microscopy on Huh7 hepatic cells using colloidal gold-conjugated anti-FITC antibodies to specifically label PGN (with no gold signals in control cells). The electron micrographs clearly showed that gold particles (representing PGN) localize to lysosomal structures, confirming stable spatial co-localization with this organelle (Figure 4F).

Collectively, these data indicate that PGN is taken up by cells and localizes to lysosomes, suggesting a possible role in regulating lysosomal functions.

### PGN promotes lysosomal acidification and cathepsin activity to extend lifespan

To examine how PGN affects lysosomal function in *C. elegans*, we first assessed lysosomal dynamics using the NUC-1::mCherry reporter. Kendall’s Tau-b correlation analysis of time-lapse images taken 15 seconds apart revealed a significantly lower correlation coefficient in PGN-supplemented animals at Adult+5day and Adult+10d (Figure 5A, Figure S6A-B), indicating that PGN treatment enhances lysosomal motility and dynamics. We next directly examined lysosomal acidification using the pH-sensitive fluorescent reporter NUC-1::pHTomato (Sun et al., 2020), in which higher fluorescence intensity reflects higher (less acidic) luminal pH (Li and Tsien, 2012) (Figure 5B). PGN-treated animals displayed significantly lower NUC-1::pHTomato fluorescence intensity (Figure 5C, Figure S6C-D), demonstrating that PGN promotes lysosomal acidification.

**Figure 5.**
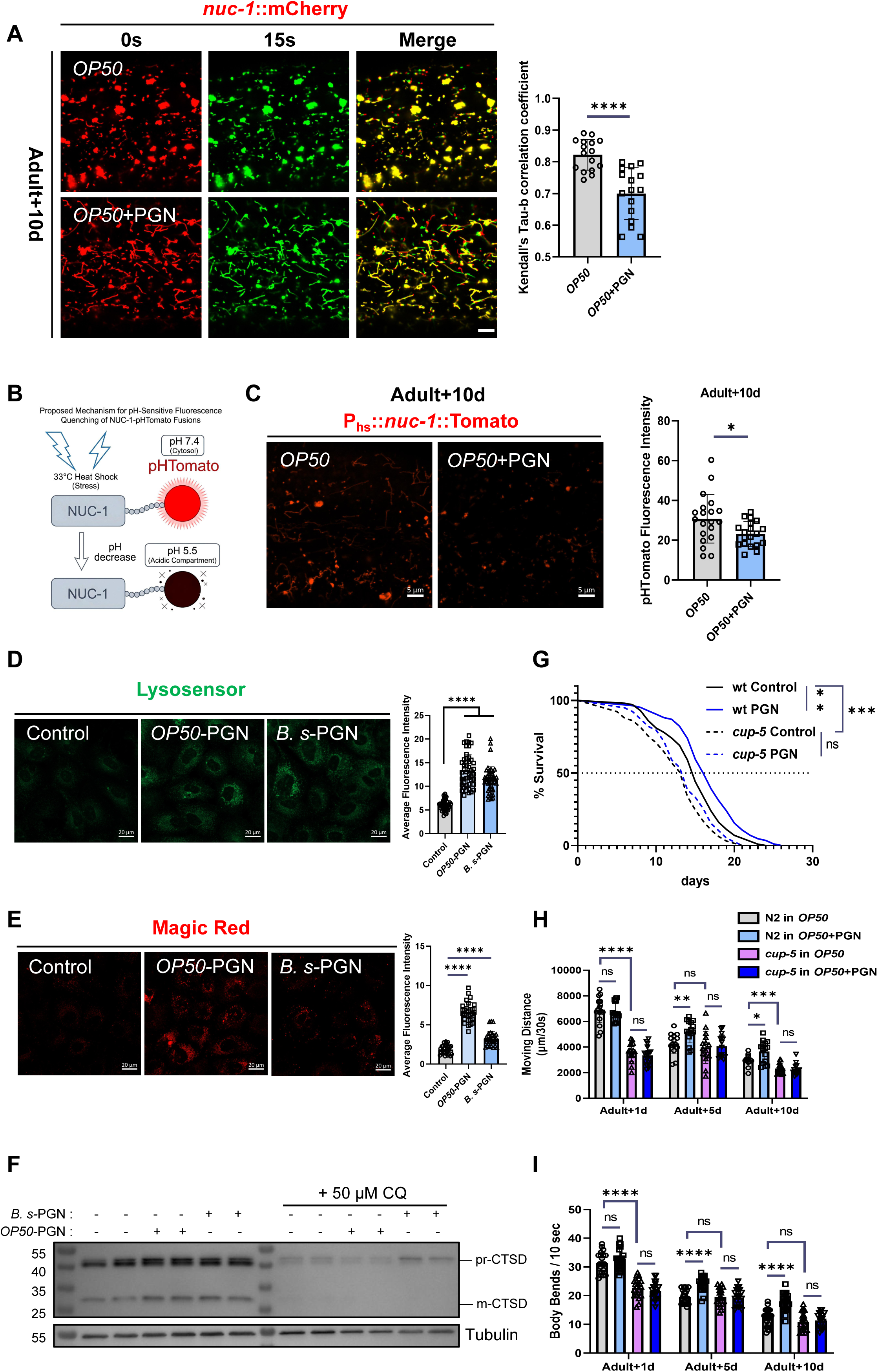
PGN promotes lysosomal activity to extend lifespan; lysosomal function is required for PGN-mediated longevity. **(A)** Representative confocal images of NUC-1::mCHERRY lysosomes at 0 s and 15 s, and merged image, in Adult+10d worms on OP50 or OP50+PGN. Bar graph shows Kendall’s Tau-b correlation coefficients (lower = higher lysosomal dynamics). Scale bar, 5 μm. Data are mean ± SD. ****P < 0.0001 (by unpaired two-tailed Student’s t-tests). n ≥ 10 worms per group; N= 3 independent biological replicates. **(B)** Schematic illustrating the NUC-1::pHTomato lysosomal pH reporter: pHTomato fluorescence increases at higher pH (lower acidity); lysosomal acidification is reflected by decreased fluorescence after pH decrease. **(C)** Representative confocal images and quantification of pHTomato fluorescence intensity per lysosome in Adult+10d worms on OP50 or OP50+PGN. Scale bar, 5 μm. Data are mean ± SD. *P < 0.05 (by unpaired two-tailed Student’s t-tests). n ≥ 20 worms per group; N= 3 independent biological replicates. **(D)** LysoSensor Green staining and quantification in Huh7 cells treated with control, OP50-PGN, or B.s-PGN. Scale bar, 20 μm. Data are mean ± SD. ****P < 0.0001 (by unpaired two-tailed Student’s t-tests). n ≥ 40 cells per replicate; N=3 independent biological replicates. **(E)** Magic Red cathepsin activity assay and quantification in control, OP50-PGN, and B.s-PGN treated cells. Scale bar, 20 μm. Data are mean ± SD. ****P < 0.0001, *P < 0.05 (by unpaired two-tailed Student’s t-tests). n ≥ 30 cells per replicate; N=3 independent biological replicates. **(F)** Western blot showing CTSD pro-form (35 kDa) and active form (25 kDa) in Adult+10d worms treated with B.s-PGN, OP50-PGN, or OP50-PGN + 50 μM chloroquine (CQ). Tubulin serves as loading control. **(G)** Survival curves of N2 wild-type and *cup-5* mutant worms on OP50 or OP50+PGN (n ≥ 100 worms per replicate;N = 3 replicates). *P < 0.05, ***P < 0.001; ns, not significant (by log-rank (Mantel-Cox) test). **(H, I)** Moving distance (H) and body bends (I) of N2 and *cup-5* worms on OP50 or OP50+PGN at Adult+1d, +5d, +10d. Data are mean ± SD. ****P < 0.0001; ns, not significant (by unpaired two-tailed Student’s t-tests). See also Figure S6 and Table S1.

In mammalian cells, treatment with both OP50-derived PGN and B.s-PGN significantly increased LysoSensor Green fluorescence, confirming enhanced lysosomal acidification (Figure 5D). Magic Red cathepsin B substrate assays demonstrated that PGN treatment markedly elevated cathepsin activity (Figure 5E), indicating that PGN evaluated lysosomal degradation activity. Western blot analysis showed that PGN increased the ratio of the mature form (m-CTSD) to the proenzyme form (pro-CTSD) of cathepsin D (CTSD), and this effect was abolished by the lysosomal alkalizing agent chloroquine (CQ, 50 μM), further confirming that PGN acts through lysosomal acidification (Figure 5F).

To determine whether lysosomal function is required for PGN-mediated lifespan extension, we tested PGN supplementation in *cup-5* mutant worms, which have severely impaired lysosomal function (Hersh et al., 2002). While PGN significantly extended lifespan in wild-type worms, it failed to do so in *cup-5* mutants (Figure 5G). Similarly, the beneficial effects of PGN on locomotor activity (Figure 5H) and body bending frequency (Figure 5I) were abolished in *cup-5* mutants. These results demonstrate that functional lysosomes are essential for the longevity-promoting effects of PGN.

Collectively, our data show that PGN extends lifespan and healthspan by enhancing lysosomal acidification and function, and this pro-longevity effect strictly requires intact lysosomes.

### PGN directly binds V-ATPase subunits and enhances V-ATPase ATPase activity

To identify the molecular target through which PGN activates lysosomal acidification, we re-analyzed the PGN pulldown proteomics data with a focus on V-ATPase subunits—the critical proton pump responsible for lysosomal acidification. In the *C. elegans* B.s-PGN interactome (Figure 4C, Table S3), multiple subunits of the V1 domain were recovered, with VHA-13 (human homolog: ATP6V1A) being the most abundantly represented (18 unique peptides), followed by VHA-12 (human homolog: ATP6V1B2, 7 peptides) and VHA-8 (human homolog: ATP6V1E1, 7 peptides) (Figure 6A; complex structure illustrated in Figure 6B).

**Figure 6.**
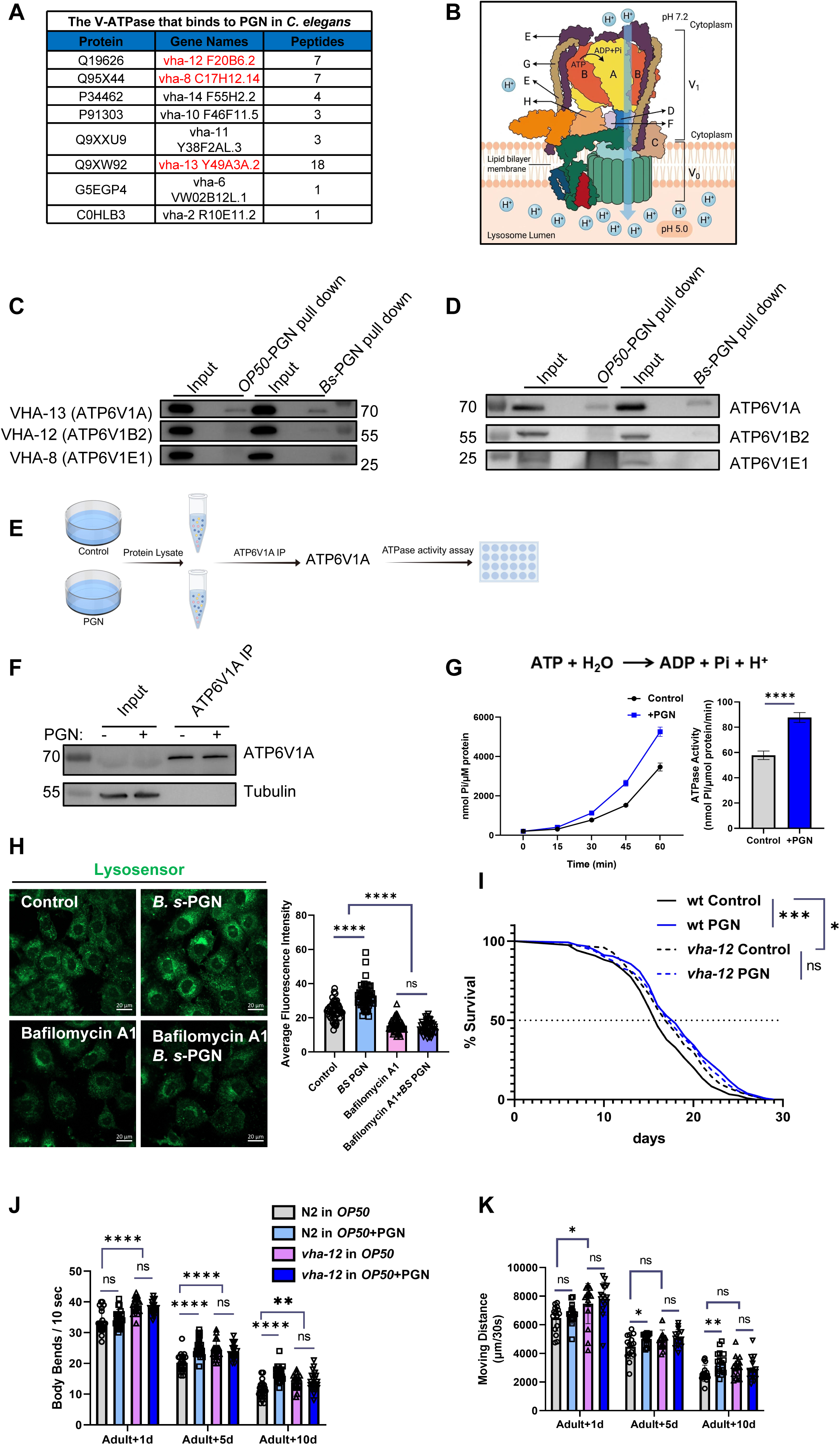
PGN directly binds V-ATPase subunits and enhances V-ATPase activity. (A) Table of V-ATPase subunits identified by PGN co-precipitation mass spectrometry in *C. elegans*, showing protein ID, gene name, and number of unique peptides. VHA-13/ATP6V1A shows the highest enrichment (18 peptides). (B) Structural schematic of the V-ATPase complex showing V_1_ (peripheral, cytoplasmic) and V_0_ (membrane-embedded) domains with labeled subunits. **(C, D)** Co-precipitation western blots confirming interaction of VHA-13/ATP6V1A, VHA-12/ATP6V1B2, and VHA-8/ATP6V1E1 with both OP50-PGN and B.s-PGN, from *C. elegans* (C) and Huh7 hepatic cell(D) lysates. (E) Schematic of the protocol for assessing V-ATPase activity in Cells treated with PGN. (F) Reciprocal immunoprecipitation (anti-ATP6V1A) from PGN-treated or control cells confirms co-precipitation of ATP6V1A. (G) V-ATPase ATPase activity measured as P_i_ release (nmol/μM protein) over 60 min from immunoprecipitated V-ATPase from control and +PGN cells. Bar graph shows mean ATPase activity (Data are mean ± SD). ****P < 0.0001 (by unpaired two-tailed Student’s t-tests). N= 3 independent biological replicates. (H) LysoSensor Green images and quantification in Huh7 cells treated with control, B.s-PGN, bafilomycin A1 (BafA1), or BafA1+B.s-PGN. Scale bar, 20 μm. Data are mean ± SD. ****P < 0.0001; ns, not significant (by unpaired two-tailed Student’s t-tests). n ≥ 50 cells per replicate; N= 3 independent biological replicates. (I) Survival curves of N2 and *vha-12(ok821)* mutant worms on OP50 or OP50+PGN (n ≥ 100 worms per replicate). *P < 0.05, ***P < 0.001; ns, not significant (by log-rank (Mantel-Cox) test). **(J, K)** Body bends (J) and moving distance (K) of N2 and *vha-12* mutants at Adult+1d, +5d, and +10d. Data are mean ± SD. **P < 0.01, ****P < 0.0001; ns, not significant (by unpaired two-tailed Student’s t-tests). n ≥ 15 worms per replicate; N=3 independent biological replicates. See also Figure S7 and Table S1.

We validated these interactions via co-precipitation followed by Western blotting, using OP50-PGN and B.s-PGN as baits in both *C. elegans* total lysates (Figure 6C) and Huh7 hepatic cell lysates (Figure 6D). The results confirmed that PGN binds to VHA-12 and VHA-13 in *C. elegans*, but binds strongly to ATP6V1 in mammalian cell. Furthermore, cell-free pull-down assays with recombinant purified ATP6V1A demonstrated that PGN binds directly to this subunit, excluding indirect bridging by adaptor proteins (Figure S7A-B).

Given that ATP6V1A is the catalytic ATPase V1 subunit that drives proton pumping through ATP hydrolysis (Figure 6B), we hypothesized that PGN binding regulates its enzymatic activity. To assess the effects of PGN on V-ATPase, we immunoprecipitated the V1 complex (ATP6V1A) from BV2 cells treated with or without PGN and measured its ATP-hydrolyzing activity (Figure 6E–F). Over a 60-min time course, samples from PGN-stimulated cells exhibited substantially higher ATPase activity than control preparations (Figure 6G), indicating that PGN engagement enhances catalytic turnover. To confirm that this enhanced V-ATPase activity is causally responsible for PGN-induced lysosomal acidification, we pretreated Huh7 cells with bafilomycin A1 (BafA1), a specific V-ATPase inhibitor (Wang et al., 2021), before adding B.s-PGN. BafA1 completely abolished the PGN-dependent increase in LysoSensor fluorescence (Figure 6H), indicating that PGN acidifies lysosomes exclusively through V-ATPase. Finally, genetic epistasis in *C. elegans* using *vha-12(ok821)* mutant revealed that PGN supplementation failed to extend lifespan (Figure 6I) or improve healthspan metrics (Figures 6J, 6K) in V-ATPase-compromised background, despite modest baseline longevity in these mutants by inducing lysosomal surveillance response (Li et al., 2025). Together, these results establish V-ATPase as the direct molecular target through which PGN activates lysosomal function to promote longevity.

### Long-term PGN supplementation maintains lysosomal homeostasis and reduces cellular senescence in multiple tissues

To investigate the tissue-level anti- aging effects of long-term PGN supplementation in mammals, we administered PGN in the diet starting at 3 months of age (early life stage) and continued the intervention until tissue collection at 20 months (Figure 7A). We first assessed lysosomal function: primary hepatocytes were isolated and stained with LysoSensor Green, and the results showed that the PGN- supplemented group exhibited significantly higher lysosomal acidification fluorescence intensity than the control group (Figure 7B), indicating that chronic PGN treatment preserves lysosomal acidification capacity during aging. Transmission electron microscopy of brain and skeletal muscle tissues revealed that PGN-treated mice had a significantly lower proportion of lysosomes with disrupted membranes in both tissues, with better- preserved structural integrity (Figure 7C-D).

**Figure 7.**
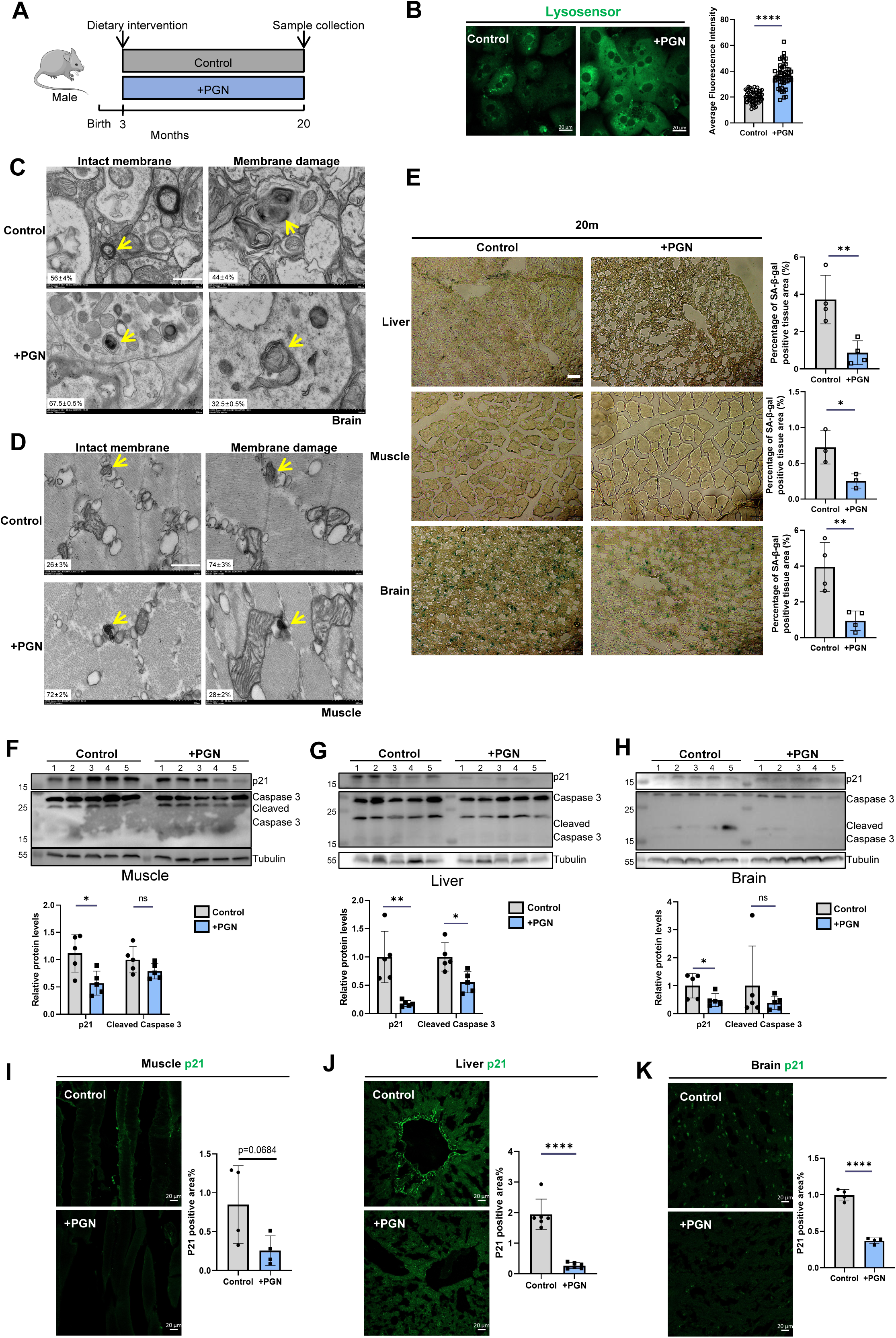
Long-term PGN supplementation in mice maintains lysosomal homeostasis, preserves tissue integrity, and reduces cellular senescence across multiple organs. **(A)** Experimental schematic: male C57BL/6J mice received PGN-supplemented or control chow from 3 months of age; tissues were collected at 20 months. **(B)** Representative LysoSensor Green images and quantification of lysosomal fluorescence intensity in primary hepatocytes from control and +PGN mice at 20 months. Scale bar, 20 μm. Data are mean ± SD. ****P < 0.0001 (by unpaired two-tailed Student’s t-tests). n ≥ 50 cells per replicate; N= 3 independent biological replicates. **(C, D)** Transmission electron micrographs of brain (C) and skeletal muscle (D) lysosomes in control and +PGN mice. Percentages indicate proportions of intact versus membrane-damaged lysosomes per field. Yellow arrows indicate lysosomal structures in muscle tissue. **(E)** Senescence-associated β-galactosidase (SA-β-gal) staining of liver (upper), skeletal muscle (middle), and brain (lower) sections from 20-month-old control and +PGN mice. Bar graphs show percentage of SA-β-gal-positive tissue area. **P < 0.01, *P < 0.05 (by unpaired two-tailed Student’s t-tests). n≥ 3 mice/group. **(F–H)** Western blots for p21, Caspase-3, cleaved Caspase-3, and tubulin in skeletal muscle (F), liver (G), and brain (H) from control and +PGN mice (n = 5 per group). Bar graphs show quantified relative protein levels.Data are mean ± SD. *P < 0.05, **P < 0.01; ns, not significant (by unpaired two-tailed Student’s t-tests). **(I–K)** Immunofluorescence images and quantification of p21-positive area in skeletal muscle (I; P = 0.0684), liver (J; ****P < 0.0001), and brain (K; ****P < 0.0001) from control and +PGN mice. Scale bar, 20 μm. Data are mean ± SD. ****P < 0.0001 (by unpaired two-tailed Student’s t-tests). n≥ 4 mice per group.

**Figure 8.**
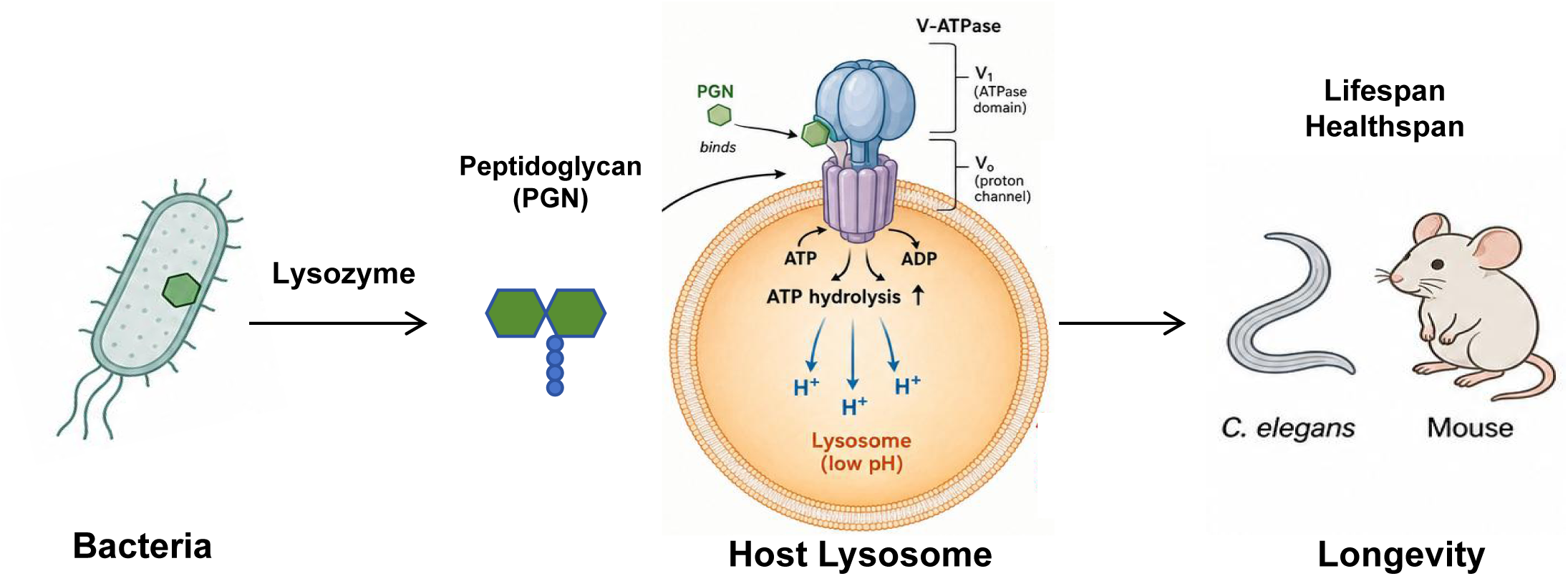
A schematic model of this study. Microbiota-derived peptidoglycan (PGN) acts as a systemic longevity signal, extending lifespan in both *C. elegans* and mice through direct binding to V-ATPase subunits and subsequent lysosomal acidification. This study indicates that the age-related decline in intestinal lysozyme activity impairs bacterial cell wall degradation, consequently diminishing PGN bioavailability. Importantly, exogenous PGN supplementation compensates for this deficit by restoring lysosomal function, thereby prolonging both healthspan and overall lifespan. Collectively, these findings establish a new paradigm that bridges microbiome metabolism, lysosomal homeostasis, and systemic aging.

We further examined senescence-related molecular markers, including the β-galactosidase (SA-β-gal) (Debacq-Chainiaux et al., 2009), cell-cycle inhibitor p21 (Hara et al., 1996) and the pro-apoptotic inflammatory molecule cleaved caspase-3 (Zhang et al., 2002). SA-β-gal staining of liver, brain, and skeletal muscle demonstrated markedly reduced positive signals in the PGN-treated group across all three tissues (Figure 7E). Western blot analysis showed that p21 protein expression was downregulated to varying degrees in the liver, brain, and skeletal muscle of PGN-fed mice (Figures 7F-H), whereas cleaved caspase-3 levels were significantly reduced only in the liver tissue (Figures 7G). Immunofluorescence staining for p21 confirmed these findings, with significantly fewer p21-positive cells in all tissues examined in the PGN-treated group compared with controls (Figures 7I-K).

Collectively, these data demonstrate that PGN supplementation maintains lysosomal homeostasis across multiple tissues during aging and reduces the accumulation of senescent cells, thereby contributing to organism-wide anti-aging effects.

## Discussion

In this study, we have identified bacterial peptidoglycan as a previously unrecognized microbiota-derived longevity signal that acts by directly engaging the V-ATPase to enhance lysosomal acidification. These findings establish a new gut microbiota–lysosome–longevity axis and identify PGN as a naturally occurring pro-longevity compound with broad therapeutic potential for promoting healthy aging.

Our findings redefine the known biology of PGN and its role in aging. The classical role of PGN as a pathogen-associated molecular pattern (PAMP) triggering innate immune signaling through NOD1/NOD2, TLR2/CD14, and PGRP receptors to activate NF-κB and MAPK pathways has been extensively studied (Dziarski, 2003; Girardin et al., 2003; Wolf and Underhill, 2018). More recently, Tian and Han (2022) (Tian and Han, 2022) demonstrated that PGN muropeptide fragments act as direct agonists of the mitochondrial F-type ATP synthase in *C. elegans* intestinal cells, supporting mitochondrial homeostasis by repressing oxidative stress; this finding was subsequently extended to mammals (Tian et al., 2024). Our data now reveal that PGN—targeting the lysosomal V-ATPase rather than the mitochondrial F-ATPase—constitutes a lysosomal activity-enhancing signal. This conceptual expansion is significant because it suggests that host cells have co-opted the structural component of their microbiota not only as a immunalsignal or a mitochondrial co-factor, but also as an organellar maintenance molecule: a “microbiota-derived lysosomal tonic” whose age-dependent depletion predisposes to lysosomal dysfunction. The mechanistic parallel between PGN’s engagement of the F-ATPase and the V-ATPase is compelling: both ATPases belong to the rotary-mechanism superfamily; both perform proton pumping coupled to nucleotide hydrolysis or synthesis; and both are now demonstrated to interact directly with PGN as natural agonists. This reveals an unexpected evolutionary depth to the molecular dialogue between bacterial cell wall chemistry and eukaryotic bioenergetic machinery.

This work fills a critical gap in the lysosome-aging field. While it has been well established that lysosomal activity declines during aging in yeast (Hashmi and Kane, 2025), nematodes (Sun et al., 2020), and mammals (Burrinha et al., 2023; Nixon, 2020), and that overexpression of V-ATPase subunits can restore acidification and extend lifespan (Ruckenstuhl et al., 2014), no naturally occurring, exogenously administrable molecule capable of directly binding V-ATPase V1 and enhancing its catalytic activity had previously been identified. PGN is such a molecule. Its direct binding to ATP6V1A/VHA-13—demonstrated here by cell-free reconstitution with purified protein—and the consequent enhancement of ATP hydrolytic activity position PGN as the first physiologically sourced V-ATPase V1 agonist. This contrasts sharply with bafilomycin A1, which is an inhibitor used only as an experimental tool. The clinical implications of identifying a food-grade, bacterially derived activator of the lysosomal proton pump are therefore considerable.

Our work also introduces a conceptually new dimension to the rapidly growing field of microbiota–aging interactions. Almost all previous studies attribute microbiota–longevity connections to either specific bacterial species composition, diffusible metabolites, or immune modulation (Caetano-Silva et al., 2024; Golomb et al., 2020; Gusarov et al., 2013; Han et al., 2017; Li et al., 2016; O’Toole and Jeffery, 2015; Thevaranjan et al., 2017). Our study takes a fundamentally different approach by focusing on bacterial structural components—specifically PGN itself—and on a non-immune effector mechanism (lysosomal V-ATPase activation). This framework reframes bacteria not merely as metabolic factories or immune stimulants, but as providers of structural biomolecular signals that directly maintain host organellar homeostasis—a function whose gradual loss during aging constitutes a new upstream driver of cellular degeneration. By showing that the age-related decline in lysozyme leads to reduced circulating PGN fragments (evidenced by decreased serum NAM in aged mice), we provide direct evidence that aging itself causes a systemic deficiency in a microbiota-derived organellar maintenance molecule.

Furthermore, our study introduces the concept of a gut–lysosome aging axis—a previously unrecognized inter-organ communication relay through which the progressive failure of intestinal digestive function directly degrades the lysosomal capacity of cells throughout the body. This differs from the well-characterized gut–brain or gut–immune axes in that its effector are a diffusible structural molecule (PGN/muropeptides) that penetrates target cells and engages their core housekeeping machinery. The evidence for systemic distribution of PGN is supported by our serum NAM mass spectrometry data, consistent with published evidence that PGN enter systemic circulation (Wheeler et al., 2023). This gut–lysosome axis places intestinal lysozyme secretion as an upstream determinant of systemic lysosomal health, suggesting that age-related Paneth cell dysfunction—well documented in humans and mice (Pentinmikko et al., 2019; Yu et al., 2020; Zhang et al., 2025)—may be a critical and underappreciated proximal cause of the lysosomal aging observed in distant tissues including liver, brain, and muscle.

In summary, our findings establish a new molecular node in the biology of aging that directly connects gut microbial cell wall to intracellular organellar function. We demonstrate that intervening at this node—by supplementing a structurally simple bacterial molecule (PGN)—prolongs healthy lifespan in two evolutionarily distant organisms. Future work will be required to elucidate the structural basis of the PGN–V-ATPase interaction, map the tissue specificity of this effect, and translate these findings to humans. Nonetheless, our findings represent a significant conceptual advance in our understanding of how the gut microbiota’s cell wall contributes to cellular aging, while also highlighting a promising avenue for the development of novel anti-aging therapeutics.

### Limitation of the study

Several limitations and open questions merit future investigation. First, the precise structural requirements for PGN-ATP6V1A interaction remain unknown: whether intact PGN polymers, specific muropeptide sizes, or particular peptide-stem configurations are the active species requires biochemical and structural characterization, potentially using cryo-EM of the PGN-V-ATPase complex. Second, the tissue specificity of PGN uptake and lysosomal effects across organs (liver, muscle, brain, kidney) deserves systematic mapping, as tissue-specific lysozyme content and PGN bioavailability may produce heterogeneous effects. Third, whether oral PGN supplementation in humans produces comparable circulating NAM increases and lysosomal functional improvements is a critical outstanding question for translation. Fourth, the relationship between the PGN-V-ATPase axis and established longevity pathways (IIS, mTOR, AMPK) warrants exploration, as PGN-driven lysosomal activation may modulate mTORC1 or AMPK signaling through the lysosomal surface. These future directions notwithstanding, our findings define a new molecular node in the biology of aging that connects gut microbial ecology to intracellular organellar function through a specific, targetable molecular interaction, and demonstrate that intervening at this node prolongs healthy lifespan in two evolutionarily distant organisms.

## Resource Availability

### Lead Contact

Further information and requests for resources and reagents should be directed to and will be fulfilled by the lead contact, Bin Qi.

### Materials availability

All unique reagents generated in this study are available from the Lead Contact upon request.

### Data and code availability

All data reported in this paper are available from the lead contact upon request.

Any additional information required to reanalyze the data reported in this paper is available from the lead contact upon request.

## Acknowledgements

We thank the Caenorhabditis Genetics Center (CGC; funded by NIH P40 OD010440) for providing worm strains, and Dr. Xiaochen Wang and Dr. Chongling Yang for sharing lysosomal reporter strains. For helpful discussions, we acknowledge Dr. Mei Yang, Dr. Zhouliang Yu, Dr. Wenxiang Fu, Dr. Qingfeng Chen, Dr. Yong Tao, and Dr. Yang Yang. We also appreciate Dr. Xuemei Niu for her technical assistance with NAM measurements, and Dr. Xuna Wu of the Mass Spectrometry Facility for mass spectrometry. Finally, we thank the members of the Qi and Shan laboratories for their insightful comments and technical support.

This work was supported by the Yunnan Provincial Science and Technology Project at Southwest United Graduate School (202302AP370005 to B.Q.), National Natural Science Foundation of China (82570734 to Z.S; 32541028 to B.Q), Yunnan Revitalization Talent Support Program (C619300A086 to Z.S., K264202230211 to B.Q.).

## Author Contributions

H.F. performed *C. elegans* experiments, proteomics analyses, biochemical characterization, mouse lifespan, healthspan studies and wrote the manuscript. J.T. performed mammalian cell experiments and mouse/tissue analyses.Y.Z. performed the *C. elegans* lifespan assays and immunostaining experiments. X.W. assisted with the transmission electron microscopy. B.Q. and Z.S. conceived the project, designed experiments, supervised the work, and wrote the manuscript with input from all authors.

## STAR METHODS

### Experimental model and study participant details

#### Animals

Male C57BL/6J mice (3 months of age, ∼20 g body weight), female C57BL/6J mice (18 months of age, 32–38 g), and male C57BL/6J mice (18 months of age, 22–30 g) were obtained from the Yunnan University Animal Center. Animals were housed in a pathogen-free facility with a 12 h light/12 h dark cycle and provided water and standard chow ad libitum at Yunnan University Animal Center (Latitude: 24°49’45.220” N; Longitude: 102°51’2.470” E; Elevation: 1,938 meters). All animal procedures were approved by the Institutional Animal Care and Use Committee of Yunnan University.

#### *C. elegans* strains and maintenance

Unless otherwise noted, worms were maintained on standard NGM agar plates seeded with *E. coli* OP50 at 20°C.

The following strains were obtained from the CGC:

N2 (wild type);

RW1596: *myo-3(st386); stEx30*;

HZ108: *cup-5(bp510)*;

RB938: *vha-12(ok821)*.

The following strains were obtained from the Dr.Xiaochen Wang laboratory:

XW5399: P*(ced-1)*NUC-1::CHERRY(qxIs257) (Sun et al., 2020);

XW19180: P*(hs)*NUC-1::pHTomato(qxIs750) (Sun et al., 2020).

Lysozyme overexpression transgenic strains were generated in this study.

YUN891: *lys-1 o/e; lys-2 o/e; lys-8 o/e(ylfEx442)*

#### Cell lines

Huh7, AML-12 and BV2 cells were maintained in DMEM (VivaCell C3113-0500) supplemented with 10% fetal bovine serum (FBS; VivaCell C04001-500) and penicillin–streptomycin (VivaCell C3421-0100) at 37°C. Primary hepatocytes isolated from C57BL/6J mouse liver were cultured in Williams’ E medium (Thermo Fisher Scientific 12551032) supplemented with 10% FBS and penicillin–streptomycin at 37°C. Cells were allowed to attach fully before any experimental treatment.

#### Bacterial strains

*E. coli* OP50, *E. coli* OP50-GFP, and *Bacillus subtilis* were grown overnight at 37°C in LB medium.

## METHOD DETAILS

### Generation of transgene strains

To construct the *C. elegans* plasmid for expression of P*lys-1*::LYS-1::mCherry, P*lys-2*::LYS-2::mCherry and P*lys8*::LYS-8::mCherry, the corresponding promoter fragments (2004 bp for *lys-1*, 1775 bp for *lys-2*, and 2001 bp for *lys-8*) along with the respective genomic DNA regions (1088 bp, 1074 bp, and 1362 bp, respectively) were amplified and cloned into the pPD49.26-mCherry vector. DNA plasmid mixture containing P*lys-1*::LYS-1::mCherry (20ng/µl), P*lys-2*::LYS-2::mCherry (20ng/µl), P*lys8*::LYS-8::mCherry (20ng/µl) and P*odr-1::*GFP(50ng/µl) was injected into the gonads of adult wild type N2.

### PGN extraction and preparation

PGN was extracted from *Escherichia coli* OP50 or *Bacillus subtilis*. For worm experiments, 60 mL of an overnight bacterial culture in LB medium was pelleted by centrifugation (8,000 × g, 5 min) and resuspended in 6 mL of 1 M NaCl (prepared by dissolving 1.168 g NaCl in 20 mL ddH₂O). The suspension was boiled at 100°C for 30 min, washed three times with ddH₂O, and sonicated on ice for 1 h. After centrifugation (10,000 × g, 5 min), the pellet was subjected to sequential enzymatic digestion in 14.67 mL of buffer (pH 6.8 Tris-HCl) containing DNase (5 mg/mL, 150 μL) and RNase (5 mg/mL, 180 μL) at 37°C for 1 h, followed by incubation with trypsin (50 μg/mL; Sangon, A100260-0050) at 37°C for an additional 1 h. The sample was then washed three times with ddH₂O, re-boiled at 100°C for 5 min, and the purified PGN was collected by centrifugation (8,000 × g, 5 min) and and quantified by weight before use.

For mouse experiments, 1 L of an overnight culture of B. subtilis ATCC 6051 in LB medium was centrifuged (8,000 × g, 5 min) to pellet the bacterial cells. The pellet was resuspended in 40 mL of 1 M NaCl, boiled at 100°C for 30 min, washed three times with ddH₂O, and sonicated on ice for 1 h. After centrifugation (10,000 × g, 5 min), the pellet was resuspended in 80 mL of digestion buffer (pH 6.8 Tris-HCl) containing DNase (5 mg/mL, 1,600 μL) and RNase (5 mg/mL, 1,920 μL), and incubated at 37°C for 1 h. Subsequently, trypsin (50 μg/mL; Sangon Biotech, A100260-0050) was added and the mixture was further incubated at 37°C for 1 h. Following enzymatic digestion, the sample was washed three times with ddH₂O, re-boiled at 100°C for 5 min, and the purified PGN was collected by centrifugation (8,000 × g, 5 min). PGN derived from 1 L bacterial culture was incorporated into 1 kg of standard mouse maintenance chow.

For FITC labeling, 500 mg PGN was incubated with 1 mg/mL FITC (Sigma F7250) in the dark for 1 h at room temperature. Excess FITC was removed by 3× PBS washes and labeling efficiency was verified by microscopy.

### *C. elegans* lifespan assays

Synchronized L1-stage worms were cultured on OP50-seeded NGM plates at 20°C for approximately 2 days until they reached the late L4 larval stage. Individual worms were then transferred to 30 mm NGM plates supplemented with 5-fluorodeoxyuridine (FUDR) to prevent progeny production. Each plate contained 20-30 worms, and five biological replicate plates were set up for each experimental condition. Survival was scored daily by gentle tapping on the plate or by lightly prodding the worms with the pick; worms that failed to show any spontaneous movement within 10 s were recorded as dead. Scoring continued until all worms had died. The daily number of dead worms was compiled, and survival curves were plotted and analyzed using GraphPad Prism (or GraphPad software).

### *C. elegans* healthspan assays

Synchronized L1-stage worms were cultured on OP50-seeded NGM plates at 20°C until they reached the late L4 larval stage. They were then transferred to fresh NGM plates containing the designated experimental conditions (e.g., different food sources or supplements) and maintained thereafter. Locomotor performance was assessed at three time points: day 1, day 5, and day 10 of adulthood.

#### 1) Crawling trajectory assay

For crawling assays, individual worms were picked onto OP50-seeded NGM plates and allowed to initiate spontaneous movement. After 30 s of free crawling, the worms were heat-killed by brief contact with a pre-heated pick to terminate movement and preserve the complete crawling track. The tracks were photographed under a stereomicroscope, and the total crawling distance was measured using ImageJ software. At least 15 worms were analyzed per group.

#### 2) Swimming body bend frequency assay

For swimming assays, individual worms were transferred into 1 mL of M9 buffer in a 30-mm diameter plastic dish. After a brief adaptation to the liquid environment, the number of body bends was counted manually using a hand tally counter. A body bend was defined as a directional change of the anterior body; a leftward swing was scored as 1, a rightward swing as 2, and so forth, and the total number of swings within 10 s was recorded for each worm. At least 15 worms were examined per group.

### *C. elegans* muscle integrity assay

To evaluate muscle filament integrity, the transgenic strain RW1596 (P*myo-3*::GFP::myo-3) was used, which expresses a GFPtagged myosin heavy chain A (MYO3) that labels body-wall muscle. Worms were examined at three time points: day 1, day 5, and day 10 of adulthood. At each time point, worms were anesthetized with levamisole, mounted on glass slides with coverslips, and immediately imaged using a confocal fluorescence microscope equipped with a 63× oil-immersion objective. At least 15 worms were photographed per group per time point. Muscle fibers were scored as intact if they displayed uniform fluorescence intensity, no blurring, and no visible breaks or discontinuities. The proportion of worms with intact muscle filaments was calculated for each experimental group.

### *C. elegans* intestinal permeability assay (Smurf assay)

Synchronized L1-stage worms were cultured on NGM plates seeded with OP50 *E. coli* supplemented with PGN (as per experimental design). To obtain age-matched cohorts at three time points—day 1, day 5, and day 10 of adulthood—worms were cultured in parallel and harvested on the designated days. Throughout the culture period, worms were transferred to fresh plates before the bacterial lawn was depleted to ensure ad libitum feeding and avoid starvation-induced stress.

At each time point, worms were collected by washing the plates with M9 buffer and pelleted into 1.5-mL microcentrifuge tubes. The worms were then incubated with 300 μL of 75 mg/mL FD&C Blue No. 1 disodium salt (Sigma-Aldrich, 861146) in M9 buffer at room temperature in the dark for 2–3 h. Following incubation, the worms were washed five times with M9 buffer to remove excess dye. Subsequently, worms were mounted on agarose pads and imaged under a stereomicroscope; at least 15 worms were photographed per group per time point. Intestinal barrier integrity was evaluated based on the extent of blue dye penetration through the body wall: worms were classified as having intact gut barrier (no blue dye visible in the body cavity), moderate permeability (partial dye infiltration), or severe permeability (widespread dye distribution throughout the body). The proportion of worms in each category was calculated for each experimental group.

### *C. elegans* intestinal bacterial colonization assay

The method was modified from He et al 2023 (He et al., 2023). Synchronized L1-stage worms were cultured on NGM plates seeded with OP50-GFP (a GFP-expressing E. coli OP50 strain) supplemented with PGN (as per experimental design) at 20°C. To obtain age-matched cohorts, worms were cultured in parallel and harvested at three time points: day 1, day 5, and day 10 of adulthood. Throughout the culture period, worms were transferred to fresh plates before the bacterial lawn was depleted to ensure ad libitum feeding and avoid starvation-induced stress.

At each time point, approximately 50 worms were picked from the plates, anesthetized with levamisole, and transferred to the center of LB agar plates supplemented with ampicillin. The plates were then incubated at 40°C overnight to heat-kill the worms. The following day, individual worms were carefully repositioned from the center to the periphery of the plates using a pick, taking care to minimize bacterial carryover from the central inoculation area. Worms were then imaged under a stereofluorescence microscope with identical acquisition settings (e.g., exposure time, gain, and intensity) applied across all groups to ensure comparability. Intestinal GFP fluorescence intensity, reflecting the abundance of OP50-GFP bacterial colonization in the worm gut, was quantified using ImageJ software.

### Lysosomal dynamics analysis in *C. elegans*

Lysosomal dynamics were assessed using the transgenic reporter strain XW5399 [P(ced-1)NUC-1::mCherry(qxIs257)], which expresses a lysosomal-localized mCherry fusion protein. Worms were cultured under different food conditions and examined at three time points: day 1, day 5, and day 10 of adulthood. At each time point, worms were anesthetized with levamisole, mounted on glass slides with coverslips, and immediately imaged using a laser-scanning confocal microscope equipped with a 63× oil-immersion objective. Two sequential fluorescence images of the same worm were captured at 0 s and 15 s. For each worm, the correlation between the two time-point images was quantified using ImageJ by calculating Kendall’s Tau-b correlation coefficients, which serve as an indicator of lysosomal dynamic changes: lower correlation values reflect higher lysosomal motility. At least 10 worms were analyzed per group per time point.

### NUC-1::pHTomato pH reporter assay in *C. elegans*

XW19180 [P*(hs)*NUC-1::pHTomato(qxIs750)] worms were heat-shocked at 33°C for 30 min and recovered at 20°C for 24 h before imaging. Mean pHTomato fluorescence intensity per lysosome was quantified by ImageJ (≥20 worms per group).

Lysosomal pH was assessed using the transgenic reporter strain XW19180 [P(hs)NUC-1::pHTomato(qxIs750)], which expresses a pH-sensitive fluorescent protein targeted to lysosomes. Worms were cultured under different food conditions and examined at three time points: day 1, day 5, and day 10 of adulthood. At each time point, worms were subjected to heat-shock at 33°C for 30 min to induce transgene expression, then transferred to 20°C and allowed to recover for 24 h before imaging. Following recovery, worms were anesthetized with levamisole, mounted on glass slides, and immediately imaged using a laser-scanning confocal microscope. Fluorescence images were acquired under identical settings across all groups. The mean pHTomato fluorescence intensity per lysosome was quantified using ImageJ software, with at least 20 worms analyzed per experimental group to ensure statistical reliability.

### PGN treatment of cultured cells

PGN extracted from 60 mL of bacterial culture (as described above) was resuspended in DMEM by centrifugation-based solvent exchange to generate a working stock solution. For cell treatment, 40 μL of the PGN working stock was added per 1 mL of cell culture medium; control cells received an equal volume of DMEM to exclude any solven-related effects. Cells (Huh7, AML12, BV2) were subcultured and grown for 24 h until reaching complete adherence. The cells were then treated with the PGN working solution for 20 h. After treatment, cells were harvested for subsequent analyses.

### LysoSensor Green staining

LysoSensor Green DND-189 (Thermo Fisher, L7535), a pH-dependent fluorescent probe that exhibits markedly enhanced fluorescence in acidic environments (optimal at approximately pH 5.2), was used to assess lysosomal acidification. The method was modified from Zhang et al 2026 (Zhang et al., 2026). Briefly, the culture medium was removed and the cells were washed twice with PBS to eliminate residual medium. The cells were then incubated in DMEM containing LysoSensor Green (at a concentration of 1 μL/mL DMSO) at 37°C for 1-2 min. After incubation, cells were immediately imaged using a laser-scanning confocal microscope to capture fluorescence signals. The mean fluorescence intensity was quantified using ImageJ software.

### Magic Red cathepsin B activity assay

Magic Red (Abcam, ab270772) is a membrane-permeable fluorescent substrate that penetrates into cells and organelles, where it is specifically cleaved by cathepsin B within functional lysosomes, generating red fluorescence. The dye was first reconstituted in DMSO to prepare a 250× stock solution according to the manufacturer’s instructions. This 250× stock was then diluted 1:10 with DMSO to obtain a 25× working stock. For cell staining, 20 μL of the 25× working stock was added to 480 μL of DMEM culture medium and mixed thoroughly to yield 500 μL of staining solution. The original culture medium was removed from cells (Huh7), and the staining solution was added to the cells. Cells were then incubated in the dark at 37°C for 30 min. After incubation, fluorescence images were captured using a laser-scanning confocal microscope, and the mean fluorescence intensity was quantified using ImageJ software.

### Western blot sample preparation

For *C. elegans*: Worms from a 90 mm plate were washed off with M9 buffer, collected into 1.5 mL microcentrifuge tubes, and washed several times with M9 buffer to remove residual bacteria. After centrifugation, the supernatant was discarded, and the worm pellet was resuspended in 300 μL of lysis buffer supplemented with protease inhibitors (100× PMSF and cOmplete protease inhibitor cocktail) and 6× SDS loading buffer. The mixture was heated at 100°C for 15 min to fully lyse the worms. The lysate was then centrifuged at 13,000 rpm for 5 min at 4°C, and the supernatant was collected as the sample for Western blot analysis.

For cultured cells: Cells grown in 6-well plates were washed with PBS, and 400 μL of RIPA lysis buffer containing protease inhibitors (100× PMSF and cOmplete protease inhibitor cocktail) was added to each well. Cells were detached using a cell scraper, and the lysates were transferred to 1.5 mL microcentrifuge tubes. After adding 6× SDS loading buffer, the samples were heated at 100°C for 15 min, then centrifuged at 13,000 rpm for 5 min at 4°C. The supernatant was collected for subsequent analysis. Anti-Cathepsin D (abcam, cat #ab75852, 1:3000) was used the experiments shown in Figure 5.

For mouse tissues: Snap-frozen tissues stored at −80°C were pulverized into fine powder using a mortar and pestle under liquid nitrogen. Approximately 30 mg of the tissue powder was transferred to a 1.5 mL microcentrifuge tube containing 1 mL of RIPA lysis buffer supplemented with protease inhibitors (100× PMSF and cOmplete protease inhibitor cocktail). Stainless steel beads were added, and the samples were further homogenized using a tissue homogenizer. The homogenates were centrifuged at 13,000 rpm for 5 min at 4°C, and the supernatants were collected. The supernatants were mixed with 6× SDS loading buffer and heated at 100°C for 15 min to denature proteins before storage or loading. P21 antibody (Proteintech, cat #10355-1-AP, 1:1000) and Caspase 3 antibody (Proteintech, cat #19677-1-AP, 1:3000) were used in the experiments shown in Figure 7.

### Mouse lifespan analysis

C57BL/6J mice (both male and female) were obtained from the Laboratory Animal Center of Yunnan University and housed in a specific pathogen-free (SPF) facility under controlled environmental conditions (temperature 22 ± 2°C, humidity 50-60%) with a 12-h light/12-h dark cycle. At 18 months of age, mice were randomly assigned to either the control group or the PGN dietary supplementation group (Figure 3A). All animals were provided with ad libitum access to water and standard chow, with or without PGN supplementation, throughout the study. To control for potential confounding effects on longevity, body weight was recorded weekly, and food intake was measured per cage twice per week to calculate average daily consumption per mouse (Figures S4A-S4D).

For survival monitoring, mice were inspected daily for general health status, with specific mortality checks performed every 2-5 days, and the date of each death was recorded as it occurred. Animals that met predefined humane endpoints—including >20% body weight loss, inability to access food or water, severe lethargy, or ulcerated tumors — were humanely euthanized and censored at the time of intervention. Survival data were plotted as Kaplan-Meier curves, and statistical differences between the control and PGN-treated groups were assessed using the two-sided log-rank (Mantel-Cox) test.

### Healthspan and Behavioral Functional Assessments in mice

To evaluate the impact of PGN on healthspan, a comprehensive battery of functional tests was performed on both male and female C57BL/6J mice following dietary PGN supplementation. Male mice began PGN treatment at 18 months of age and were assessed at 25 months of age after 7 months of treatment. Female mice were initiated on the same regimen at 18 months of age but were evaluated at 30 months of age following 12 months of treatment, reflecting the sex-specific differences in aging trajectory. All behavioral and physiological assays were conducted at the Laboratory Animal Center of Yunnan University, with experimenters blinded to treatment groups throughout data acquisition and analysis.

#### 1) Metabolic cage phenotyping

Spontaneous locomotor activity and metabolic parameters were monitored using the TSE-PhenoMaster fully automated metabolic phenotyping system (TSE Systems, Germany). Mice were individually housed in metabolic cages with corncob bedding and provided ad libitum access to food and water. Following a 1 h acclimatization period to the novel environment, locomotor activity and average movement speed were continuously recorded for 48 h. Data from both groups (n≥8 per group) were analyzed for total activity counts and mean velocity across the entire monitoring cycle.

#### 2) Treadmill exercise endurance test

Exercise capacity was assessed using a small-animal treadmill (XR-PT-10B; Shanghai Xinruan Information Technology Co., Ltd., China). The apparatus was set to a 5°incline with an electrical shock stimulus intensity of 0.3 A applied to the grid floor. Mice underwent a 1 h environmental acclimatization period prior to testing. The protocol began with a 5 min warm-up phase at a speed of 5 m/min accompanied by the 0.3 A stimulus to habituate the animals to the apparatus. Subsequently, the treadmill speed was linearly accelerated from 5 m/min to 25 m/min over a total duration of 7,200 s (120 min). Exhaustion was defined as the point at which the mouse remained on the shock grid for 10 consecutive seconds without attempting to re-engage in running, with a maximum tolerance count set to zero. The total running distance and total running time were recorded for each animal (n≥6 per group).

#### 3) Open field test

Locomotor activity and exploratory behavior were evaluated using an open field arena (XR-XZ301; Shanghai Xinruan Information Technology Co., Ltd.) coupled with the VisuTrack animal behavior analysis software (Shanghai Xinruan). Following a 1 h habituation period to the testing room, each mouse was individually placed in the center of the open field box and allowed to move freely for a continuous monitoring period of 300 s. The total distance traveled and average movement speed were automatically tracked and analyzed (n≥10 per group).

#### 4) Grip strength test

Forelimb and all-limb (four limbs) grip strength were measured using the KW-ZL-2 grip strength testing system (Shanghai Xinruan). Mice were positioned to grasp the wire mesh grid with either their forelimbs or all four limbs, depending on the specific test. Once a firm grip was established, the experimenter applied steady horizontal traction to the mouse’s tail until the animal released its hold. The peak force (in grams) exerted at the moment of release was automatically recorded by the system. For each test, at least 10 mice per group (n≥7) were assessed, with multiple consecutive measurements taken per mouse to obtain a reliable average.

### V-ATPase activity assay

In vitro ATPase activity was measured according to a previously reported method (Rule et al., 2016). ATP6V1A protein was specifically immunoprecipitated from BV2 cell lysates using protein A/G agarose beads coupled with an anti-ATP6V1A antibody. The successful pull-down of ATP6V1A was confirmed by western blotting (WB) prior to the activity assay, and the ATP6V1A-bound beads were used as the enzyme source.

The ATPase reaction was performed in 0.5 mL microcentrifuge tubes. The reaction mixture (total volume of 300 µL) contained the following components: deionized water (to volume), 60 µL of 5× HNG buffer (100 mM HEPES pH 8.5, 65 mM NaCl, 5% glycerol), 30 µL of a 1:1 mixture of MgCl₂ (1 M, Invitrogen #AM9530G) and ATP (100 mM, Thermo Scientific #R0441), and an appropriate volume of the immunoprecipitated ATP6V1A-bound beads (quantified for total protein using a BCA assay). The tubes were incubated at 37 °C for 1 h. At each time point—0, 15, 30, 45, and 60 min—50 µL aliquots were sampled from the 300 µL reaction and immediately transferred into tubes containing 200 µL of 1× HNG buffer for dilution. The reactions were then rapidly terminated by flash-freezing in liquid nitrogen. All time-course samples were subsequently stored at −80 °C until further analysis.

To quantify the released inorganic phosphate (Pi), the collected samples were assayed using a Malachite Green Phosphate Detection Kit (Sigma, cat# MAK307). A 1:1 serial dilution of Pi standards ranging from 40 µM to 0 µM was prepared in a 96-well plate, with 50 µL of each standard per well, to generate a standard curve for Pi concentration versus absorbance. The frozen reaction samples were thawed, and 50 µL of each diluted sample was added to the 96-well plate in duplicate, alongside the Pi standards. Subsequently, 100 µL of the Pi detection reagent was added to each well, mixed gently, and incubated at room temperature for 25 min in the dark. The absorbance of each well was measured at 650 nm using a microplate reader. A standard curve was constructed by plotting the absorbance values against the corresponding Pi concentrations, and the Pi concentration in each reaction sample was calculated based on the standard curve equation. The ATPase activity was then determined from the rate of Pi release over time.

### PGN-binding protein assay

To validate PGN-interacting proteins, insoluble solid PGN was used as an affinity matrix. Total proteins (lysates from *C. elegans* or Huh7 cells; or purified proteins) were mixed with the PGN matrix, followed by incubation overnight at 4°C with gentle rotation to allow binding. After incubation, the mixture was centrifuged at 12,000 rpm for 5 min at 4°C to separate the PGN-bound complexes. The pellet was then washed three times with pre-chilled PBS, with centrifugation at 12,000 rpm, 4°C between washes; the final wash supernatant (LS) was also collected to confirm removal of non-specifically bound proteins. The washed pellet (P) represents the PGN-bound protein fraction. All three fractions—post-incubation supernatant (S), final wash supernatant (LS), and washed pellet (P)—were analyzed by Western blotting. The washed pellet (P), representing the PGN-bound protein fraction, was resuspended in SDS-loading buffer and heated at 100°C for 15 min to elute and denature the bound proteins. The sample was then centrifuged at 12,000 rpm for 5 min, and the resulting supernatant was collected for downstream analyses. The final supernatants were resolved by SDS-PAGE and subjected to either mass spectrometry for comprehensive identification of PGN-interacting proteins, or western blotting to detect specific candidate proteins such as V-ATPase subunits (ATP6V1A, ATP6V1B2, and ATP6V1E1).

### Immunofluorescence (IF) staining in Cell

Cells cultured on glass coverslips were washed twice with PBS and fixed with 4% paraformaldehyde in PBS for 10 min at room temperature. After three washes with PBS (5 min each), cells were permeabilized with 0.2% Triton X-100 in PBS for 10 min at room temperature, followed by three additional PBS washes. To block non-specific binding, coverslips were incubated with blocking buffer (5% goat serum and 3% BSA in PBS) for 1 h at room temperature. The cells were then incubated with primary antibody (DSHB anti-LAMP1, Cat#1D4B, 1:200 in blocking buffer) overnight at 4°C. Following three washes with TBST (5 min each), coverslips were incubated with secondary antibody (Invitrogen Alexa Fluor™ 568 goat anti-rat IgG (H+L), 1:800 in blocking buffer) for 1 h at room temperature in the dark. After three additional TBST washes, nuclei were counterstained with DAPI for 5 min at room temperature. Coverslips were washed three times with PBS, mounted on glass slides using anti-fade mounting medium before imaging.

### Immunofluorescence Staining in tissues

For immunofluorescence staining, fresh mouse liver tissues were fixed in 4% paraformaldehyde (PFA) for 1 h at room temperature, washed three times with phosphate-buffered saline (PBS), and cryoprotected in 30% sucrose solution at 4°C overnight. Tissues were then embedded in optimal cutting temperature (OCT) compound, and frozen sections were cut at 5 μm thickness using a cryostat. The sections were post-fixed in pre-cooled acetone at −20°C for 15 min, permeabilized with 0.2% Triton X-100 in PBS, and blocked with blocking buffer (5% normal goat serum and 0.3% bovine serum albumin in PBS) for 1 h at room temperature.

Sections were incubated overnight at 4°C with a rabbit anti-p21 polyclonal antibody (Proteintech, cat# 10355-1-AP) at a 1:400 dilution. After extensive washing with TBST (Tris-buffered saline containing 0.1% Tween 20), the sections were incubated with Alexa Fluor 488-conjugated goat anti-rabbit IgG (Jackson ImmunoResearch, cat# 111-545-003) at a 1:300 dilution for 1 h at room temperature. Nuclei were counterstained with DAPI, and sections were mounted with an anti-fade mounting medium. Fluorescence images were captured using a Zeiss LSM900 confocal microscope under identical acquisition parameters across all samples to ensure comparability.

### Senescence-Associated β-Galactosidase (SA-β-gal) Staining

Senescence-associated β-galactosidase activity was assessed in frozen liver sections using a commercially available SA-β-gal staining kit (Beyotime Biotechnology, cat# C0602) according to the manufacturer’s instructions with minor modifications. Briefly, frozen sections were air-dried and rewarmed to room temperature, then washed three times with phosphate-buffered saline (PBS) for at least 5 min each wash. Sections were completely covered with the fixation solution provided in the kit and incubated at room temperature for a minimum of 15 min. After fixation, sections were rinsed three times with PBS for at least 5 min per wash, and excess PBS was carefully aspirated. Sections were then incubated with the staining working solution (mixing components A, B, C, and X-Gal at a volume ratio of 1:1:93:5) overnight at 37°C. The next day, sections were washed with PBS and mounted with aqueous mounting medium. Bright-field images were captured under a standard light microscope. Quantification of SA-β-gal-positive areas was performed by measuring the percentage of blue-stained area relative to the total tissue section area using ImageJ software (or similar), with at least three randomly selected fields per section and three sections per animal.

### Measurement of NAM Levels

Serum NAM (N-Acetylmuramic Acid) concentrations were quantified high-performance liquid chromatography-tandem mass spectrometry (HPLC-MS/MS) analysis. For sample preparation, serum samples were diluted with methanol at a 1:3 ratio (v/v), vortex-mixed for 10 min, and centrifuged at 14,000 rpm and 4 °C. The resulting supernatants were collected, transferred to fresh centrifuge tubes, and evaporated to dryness under vacuum. The dried metabolite residues were reconstituted in methanol, vortexed for 10 min, and centrifuged again at 14,000 rpm and 4 °C. The final supernatants were subjected to HPLC-MS/MS analysis.

NAM standards were prepared in methanol to generate a calibration curve with concentrations ranging from 10 to 1280 ng/mL.

HPLC-MS/MS analysis was performed on a Q Exactive Focus LC-MS/MS system equipped with a PDA detector and an Orbitrap mass analyzer, operating in both positive and negative electrospray ionization modes. Chromatographic separation was achieved on a CAPCELL PAK C18 column (5 μm, 4.6 mm × 250 mm) maintained at 40 °C. The mobile phase consisted of 0.1% formic acid in water (A) and 0.1% formic acid in acetonitrile (B), delivered at a flow rate of 1.0 mL/min. The injection volume was 10 μL. The gradient elution program was set as follows: 10% B from 0 to 2 min; 25% B at 10 min; 50% B at 30 min; 90% B at 35 min; 95% B from 36 to 40 min; and 10% B from 40.1 to 45 min.

### Immunogold electron microscopy

Huh7 cells were cultured on Formvar-gelatin-coated coverslips. The membrane was transferred into 5% BSA solution, placed in 3 mm gold sample carriers, and vitrified using a high-pressure freezer. Freeze substitution was performed in an automated freeze substitution apparatus following the schedule: −90°C for 9 h in acetone containing 0.2% glutaraldehyde and 0.1% uranyl acetate; warmed to −30°C over 12 h in the same solution; held at −30°C for 2 h with three changes of 0.2% glutaraldehyde in acetone; warmed to −20°C over 1.5 h in 0.2% glutaraldehyde in acetone; warmed to 0°C over 3 h with progressive infiltration of HM20 Lowicryl resin (50%, 75%, 90%, and 100% in acetone), followed by three changes of 100% HM20 at 0°C. The resin was polymerized under ultraviolet light at −20°C. Cryo-ultrathin sections (∼50 nm) were cut at −110°C to −120°C and transferred to Formvar/carbon-coated copper grids.

For immunolabeling, grids were first subjected to rehydration and antigen retrieval by floating on drops of 3/8 sucrose (in PHEM buffer) for 10 min, followed by 1/8 sucrose (in PHEM buffer) for 10 min, all at 4°C. After washing with PHEM buffer and then with PB (3 × 10 min), grids were incubated in PB at 37°C for 4 × 10 min. Free aldehyde groups were quenched with 0.15% glycine in PB for 5 min, and grids were rinsed in PB (3 × 2 min). Nonspecific binding was blocked with 5% BSA or 10% goat serum in PB for 30 min. Sections were then incubated with primary anti-FITC rabbit monoclonal antibody (Thermo Fisher Scientific, Cat# 701078, 1:50 in 0.1% BSA/PB) for 1 h at room temperature (or overnight at 4°C) in the dark. After washing with 0.1% BSA/PB (6 × 2 min), grids were incubated with 6 nm gold-conjugated goat anti-rabbit IgG (H+L) secondary antibody (Jackson ImmunoResearch, Cat# 111-035-003, 1:30 in PB) for 1 h in the dark, followed by washes in PB (6 × 2 min). The immunoreaction was post-fixed with 2.5% glutaraldehyde in PB for 5 min, rinsed thoroughly with double-distilled water (6 × 2 min), and contrasted on ice with a mixture of 2% methylcellulose and 2% uranyl acetate (9:1, v/v) for 5 min. Grids were loop-dried and examined under a transmission electron microscope at an accelerating voltage of 80 kV.

## QUANTIFICATION AND STATISTICAL ANALYSIS

All data are presented as mean ± SEM. Statistical analyses were performed using GraphPad Prism. Two-group comparisons were made using unpaired two-tailed Student’s t-tests. Multiple-group comparisons were performed by one-way ANOVA with appropriate post-hoc tests. Lifespan statistics were calculated by log-rank (Mantel-Cox) test. Pearson’s or Kendall’s correlation analyses were used where specified. Statistical significance thresholds: *P < 0.05, **P < 0.01, ***P < 0.001, ****P < 0.0001; ns, not significant. Sample sizes for each experiment are reported in the figure legends. All experiments were performed at least three independent times unless otherwise noted.

## Supplemental Figure Legends

**Figure S1. Lysozyme-related genes are downregulated during aging, and their overexpression reduces intestinal bacterial colonization in aged animals. Related to Figure 1**.

**(A)** Expression levels of lysozyme genes and bacterial lysis-related genes across aging stages of *C. elegans*. Data were re-analyzed from a published transcriptomic dataset (Kong et al., 2024) and normalized to the levels at day 1 (D1). Values are presented as fold changes relative to D1.

**(B)** Expression profiles of lysozyme genes across aging stages of *C. elegans*. Data were retrieved from the Aging Atlas (http://mengwanglab.org/atlas) (Gao et al., 2024) and normalized to D1 levels. Values are shown as fold changes relative to D1.

**(C)** Intestinal OP50-GFP colonization levels in wild-type (N2) and transgenic worms co-overexpressing *lys-1, lys-2,* and *lys-8* (*lys-1/2/8* OE) at the indicated time points during aging. At least 20 animals were scored per group. Data are presented as mean ± SD. *p< 0.05, **p< 0.01 (unpaired two-tailed Student’s t-test).

**Figure S2. Measurement of NAM (N-Acetylmuramic Acid) levels. Related to Figure 1**.

**(A)** Schematic diagram of the peptidoglycan (PGN) molecular network structure. Peptidoglycan is composed of cross-linked polymers of N-acetylmuramic acid (NAM) and N-acetylglucosamine (NAG). The cleavage sites of lysozyme are indicated in the diagram.

**(B)** Chromatographic analysis using commercial NAM (N-Acetylmuramic Acid) standards. The characteristic peak corresponding to the 160 ng/mL standard is indicated by an arrow. Chromatograms of NAM standards across a concentration gradient ranging from 10 to 1280 ng/mL are shown, and a standard curve was generated based on the chromatographic signals.

**(C)** Serum samples collected from 2-month-old and 18-month-old male mice were subjected to chromatographic analysis. Representative chromatograms showing NAM-specific peaks from both groups are presented.

**Figure S3. PGN supplementation ameliorates age-related functional decline in *C. elegans*. Related to Figure 2**.

**(A)** Schematic diagram of the experimental paradigm for PGN feeding. L4-stage animals cultured on OP50 plates were transferred to either standard OP50 plates or OP50 plates supplemented with PGN for aging assays. Behavioral assessments were conducted at three time points: adult day 1 (adult+1d), day 5 (adult+5d), and day 10 (adult+10d). Worms were maintained under ad libitum feeding conditions throughout the experiment.

**(B)** Schematic diagram of the locomotion assay. Worm locomotion was recorded immediately after assay initiation (a point). After 30 s (b point), animals were rapidly euthanized to preserve the complete crawling traces on OP50 plates. Representative crawling track images from each group are shown.

**(C)** Representative fluorescence images showing myofilament morphology in body-wall muscles of worms at the indicated ages under PGN intervention, using the P*myo-3*::GFP::myo-3 reporter strain to label muscle structural integrity (corresponding to Figure 2F).

**Figure S4. PGN supplementation improves healthspan in aged mice. Related to Figure 3**.

**(A, B)** Body weight (A) and food intake (B) of male mice during the PGN dietary intervention lifespan study. PGN supplementation was initiated at 18 months of age, and body weight and food intake were recorded weekly throughout the monitoring period until all mice reached natural death.

**(C, D)** Body weight (C) and food intake (D) of female mice during the PGN dietary intervention lifespan study. PGN supplementation was initiated at 18 months of age, and body weight and food intake were recorded weekly throughout the monitoring period until all mice reached natural death.

**Figure S5. PGN (*E. coli*)-binding proteins. Related to Figure 4**.

Bioinformatics analysis was performed based on a published dataset (Tian and Han, 2022) to annotate the subcellular localization of proteins in *C. elegans* that interact with *E. coli* OP50-derived PGN. A total of 168 PGN interacting proteins were classified into four categories: lysosomal, mitochondrial, ribosomal, and other/unclassified.

**Figure S6. PGN promotes lysosomal activity. Related to Figure 5**.

**(A, B)** Representative confocal images of NUC-1::mCHERRY lysosomes at 0 s and 15 s, and merged image, in Adult+1d(A) and Adult+5d(B) worms on OP50 or OP50+PGN. Bar graph shows Kendall’s Tau-b correlation coefficients (lower = higher lysosomal dynamics). Scale bar, 5 μm. Data are mean ± SD. ns, not significant, ****P < 0.0001 (by unpaired two-tailed Student’s t-tests). n ≥10 worms per group.

**(C, D)** Representative confocal images and quantification of pHTomato fluorescence intensity per lysosome in Adult+1d(C) and Adult+5d(D) worms on OP50 or OP50+PGN. Scale bar, 5 μm. Data are mean ± SD. *P < 0.05, **< 0.01 (by unpaired two-tailed Student’s t-tests). n ≥ 20 worms per group.

**Figure S7. PGN binds to ATP6V1A. Related to Figure 6**.

**(A, B)** Recombinant His-tagged ATP6V1A protein was expressed in E. coli and purified in vitro. The purified protein was incubated with *E. coli-*OP50-derived PGN (A) or B. subtilis-derived PGN (B) to verify their direct interaction in vitro.

All three fractions—post-incubation supernatant (S), final wash supernatant (LS), and washed pellet (P)—were analyzed by Western blotting (see more detail in methods section “PGN-binding protein assay”). The washed pellet (P) represents the PGN-bound protein fraction.

**Video Legends:**

**Video S1.** Spontaneous activity of female mice surviving to 28 months of age after PGN dietary intervention initiated at 18 months of age. Notably, the PGN-supplemented group (the right 3 cage) exhibited markedly higher locomotor activity compared with the control group (the left 3 cage). Related to Figure 3.

**Legends of Supplemental Tables:**

**Table S1.** Statistics for survival analysis in *C. elegans*, related to Figures 1C, 1D, 2C, 5G, 6I.

**Table S2.** Survival data in mouse, related to Figures 3B-3C,

**Table S3.** List of PGN-binding proteins from *C. elegans* lysates identified by pull-down assay with *B. subtilis*-derived PGN, followed by mass spectrometric analysis. Related to Figure 4C.

**Table S4.** List of PGN-binding proteins from mouse liver lysates identified by pull-down assay with *E. coli*-derived PGN, followed by mass spectrometric analysis. Data were derived from a previously published dataset (Tie et al., 2025). Related to Figure 4D.

**Table S5.** List of PGN-binding proteins from *C. elegans* lysates identified by pull-down assay with *E. coli*-derived PGN, followed by mass spectrometric analysis. Data were derived from a previously published dataset (Tian and Han, 2022). Related to Figure S5.

