## Supplemental Figures for "Bacterial Peptidoglycan Extends Lifespan by Activating Lysosomal Activity through V-ATPase Binding"

Figure S1

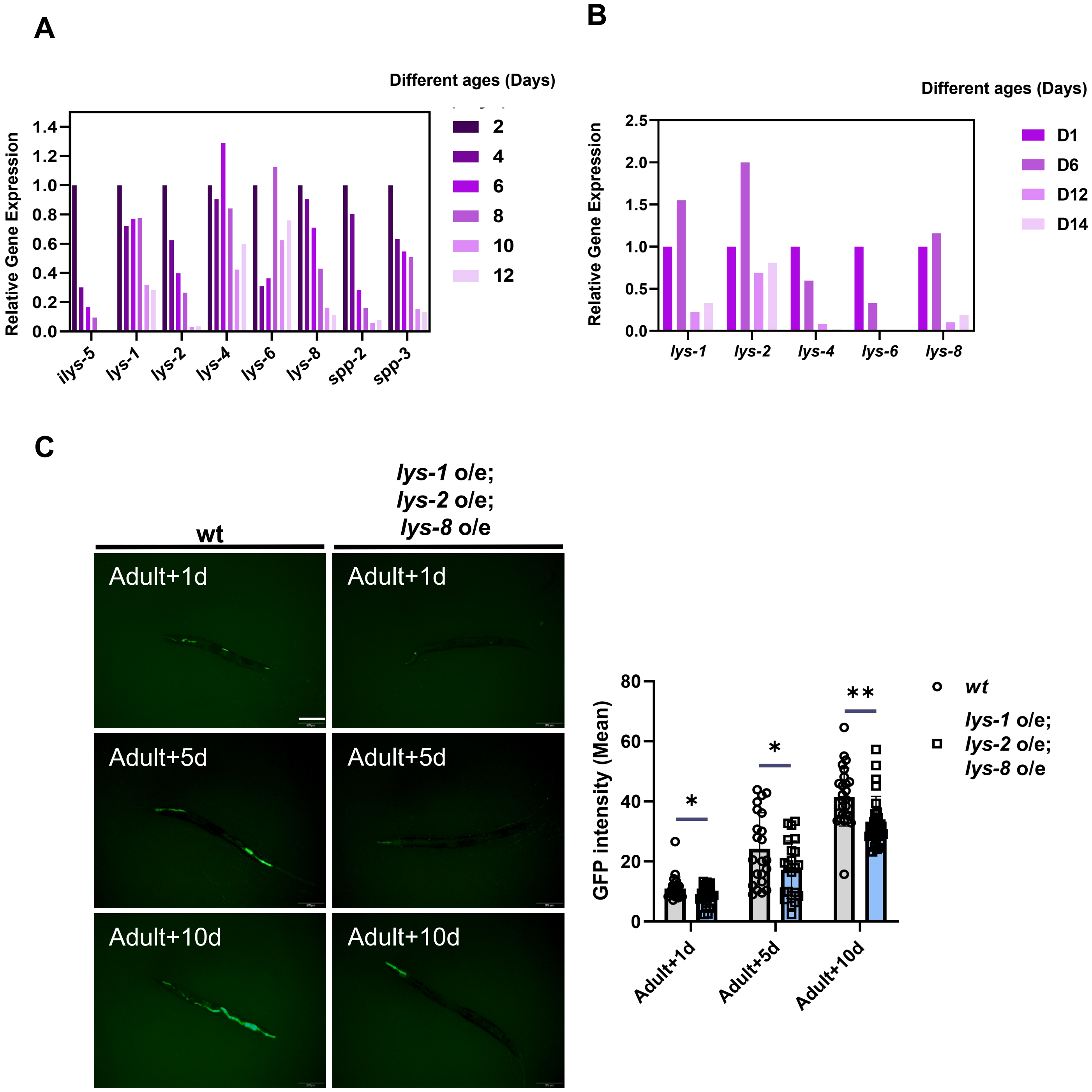


**Figure S1. Lysozyme-related genes are downregulated during aging, and their overexpression reduces intestinal bacterial colonization in aged animals. Related to Figure 1.**

**(A)** Expression levels of lysozyme genes and bacterial lysis-related genes across aging stages of *C. elegans*. Data were re-analyzed from a published transcriptomic dataset (Kong et al., 2024) and normalized to the levels at day 1 (D1). Values are presented as fold changes relative to D1.

**(B)** Expression profiles of lysozyme genes across aging stages of *C. elegans*. Data were retrieved from the Aging Atlas (http://mengwanglab.org/atlas) (Gao et al., 2024) and normalized to D1 levels. Values are shown as fold changes relative to D1.

**(C)** Intestinal OP50-GFP colonization levels in wild-type (N2) and transgenic worms co- overexpressing *lys-1, lys-2,* and *lys-8* (*lys-1/2/8* OE) at the indicated time points during aging. At least 20 animals were scored per group. Data are presented as mean ± SD. *p< 0.05, **p< 0.01 (unpaired two-tailed Student's t-test).

### Figure S2

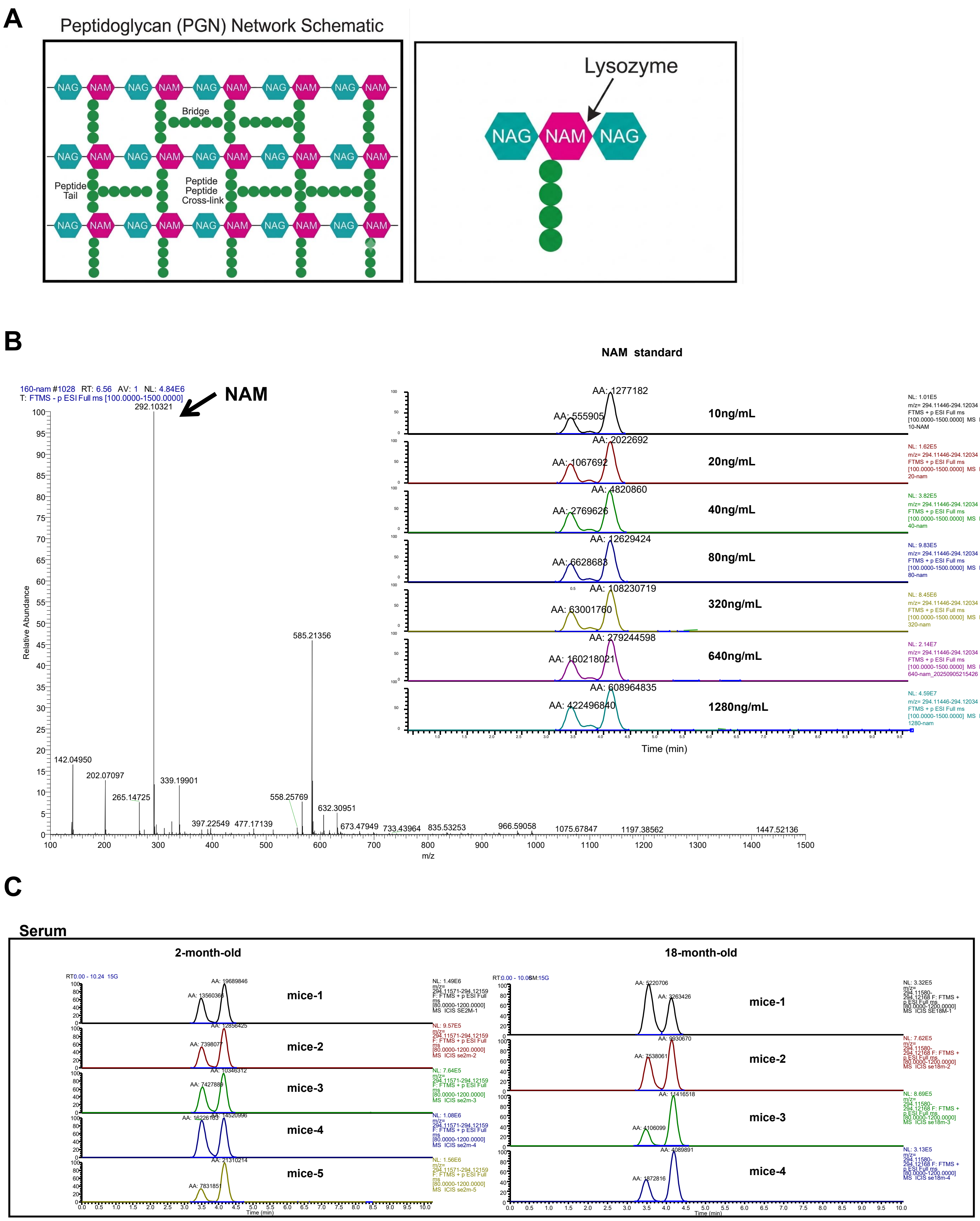


**Figure S2. Measurement of NAM (N-Acetylmuramic Acid) levels. Related to Figure 1.**

**(A)** Schematic diagram of the peptidoglycan (PGN) molecular network structure. Peptidoglycan is composed of cross-linked polymers of N-acetylmuramic acid (NAM) and N-acetylglucosamine (NAG). The cleavage sites of lysozyme are indicated in the diagram.

**(B)** Chromatographic analysis using commercial NAM (N-Acetylmuramic Acid) standards. The characteristic peak corresponding to the 160 ng/mL standard is indicated by an arrow. Chromatograms of NAM standards across a concentration gradient ranging from 10 to 1280 ng/mL are shown, and a standard curve was generated based on the chromatographic signals.

**(C)** Serum samples collected from 2-month-old and 18-month-old male mice were subjected to chromatographic analysis. Representative chromatograms showing NAM-specific peaks from both groups are presented.

**Figure S3**

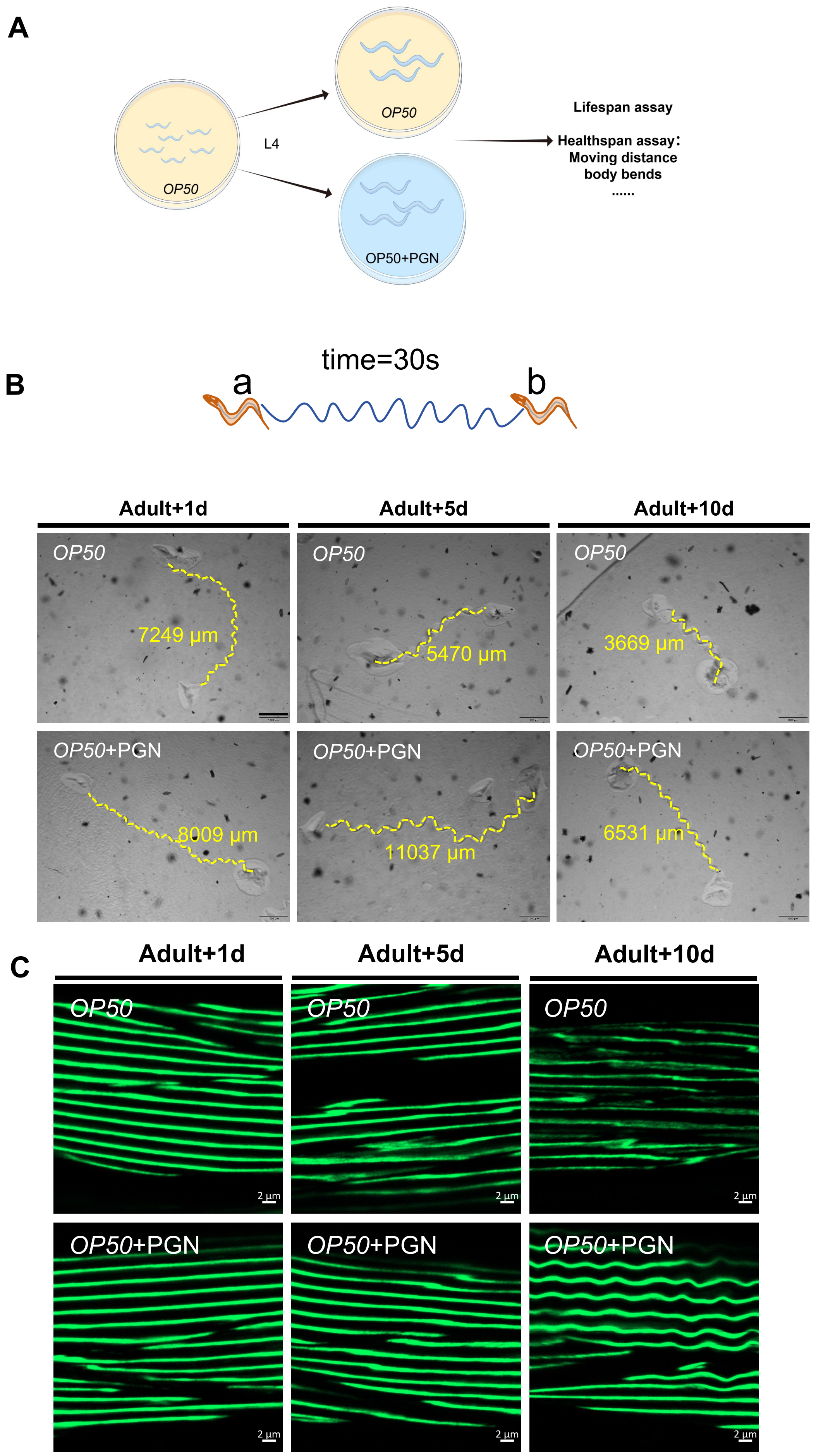


**Figure S3. PGN supplementation ameliorates age-related functional decline in *C. elegans*. Related to Figure 2.**

(A)Schematic diagram of the experimental paradigm for PGN feeding. L4-stage animals cultured on OP50 plates were transferred to either standard OP50 plates or OP50 plates supplemented with PGN for aging assays. Behavioral assessments were conducted at three time points: adult day 1 (adult+1d), day 5 (adult+5d), and day 10 (adult+10d). Worms were maintained under ad libitum feeding conditions throughout the experiment.

(B)Schematic diagram of the locomotion assay. Worm locomotion was recorded immediately after assay initiation (a point). After 30 s (b point), animals were rapidly euthanized to preserve the complete crawling traces on OP50 plates. Representative crawling track images from each group are shown.

(C)Representative fluorescence images showing myofilament morphology in body-wall muscles of worms at the indicated ages under PGN intervention, using the P*myo-3*::GFP::myo-3 reporter strain to label muscle structural integrity (corresponding to Figure 2F).

**Figure S4**

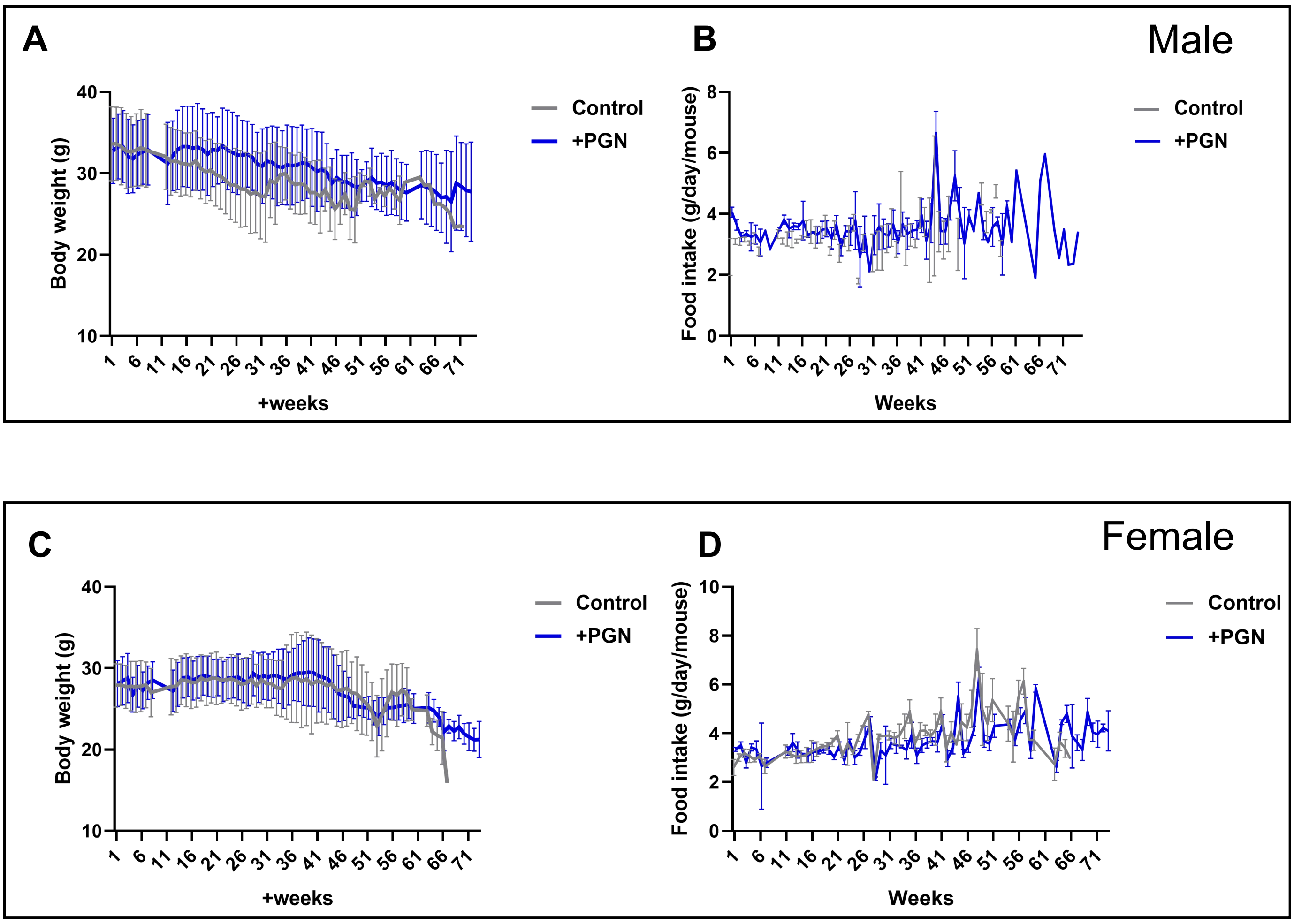


**Figure S4. PGN supplementation improves healthspan in aged mice. Related to Figure 3.**

**(A, B)** Body weight (A) and food intake (B) of male mice during the PGN dietary intervention lifespan study. PGN supplementation was initiated at 18 months of age, and body weight and food intake were recorded weekly throughout the monitoring period until all mice reached natural death.

**(C, D)** Body weight (C) and food intake (D) of female mice during the PGN dietary intervention lifespan study. PGN supplementation was initiated at 18 months of age, and body weight and food intake were recorded weekly throughout the monitoring period until all mice reached natural death.

**Figure S5**

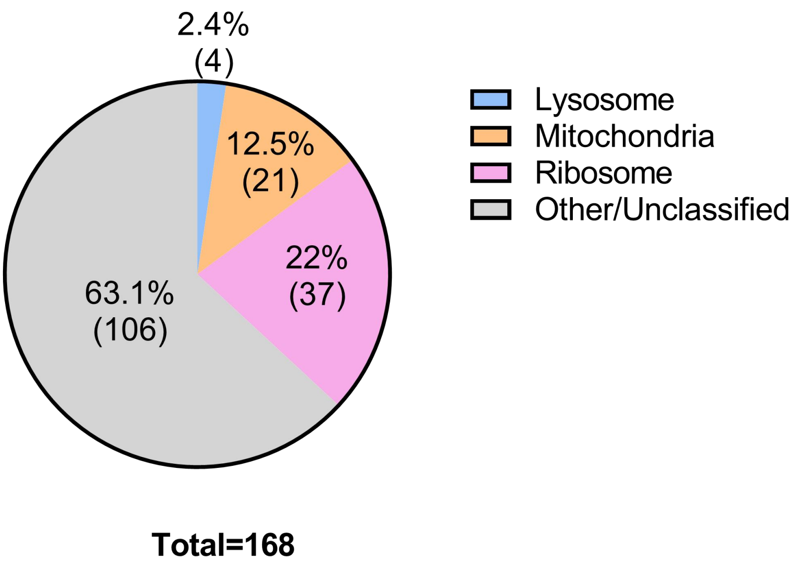


| | Number | % | Gene |
|---|---|---|---|
| **Lysosome** | 4 | 2.4 | cpl-1, ctsa-2, rme-1, vha-16 |
| **Mitochondria** | 21 | 12.5 | F23C8.5, F27D4.1, acdh-12, ant-1.1, apt-1, asb-1, atp-2, atp-4, atp-5, chch-3, dlat-1, dtmk-1, glrx-5, immt-1, kars-1, mdh-2, nuo-5, sdha-1, tim-8, tkt-1, ucr-1 |
| **Ribosome** | 37 | 22 | rla-1, rla-2, rpa-0, rpa-2, rpl-11.2, rpl-18, rpl-19, rpl-21, rpl-25.2, rpl-27, rpl-32, rpl-35, rpl-36, rpl-4, rpl-5, rpl-7A, rpl-8, rpl-9, rps-0, rps-1, rps-10, rps-12, rps-13, rps-15, rps-17, rps-18, rps-19, rps-2, rps-20, rps-21, rps-22, rps-25, rps-3, rps-5, rps-6, rps-8, ubl-1 |
| **Other/Unclassified** | 106 | 63.1 | C18B2.5, C18H9.3, F21C10.7, F21D5.1, F27D4.4, F46H5.7, F49E2.5, T09B4.5, T23E7.2, Y39B6A.33, Y75B8A.7, aars-2, act-4, act-5, alp-1, anc-1, baf-1, car-1, cey-1, cey-2, clik-1, clik-2, daf-21, dlg-1, dnc-2, ears-1, eef-1a.2, eef-2, egl-45, eif-2, eif-2alpha, fars-1, frm-1, gei-15, gfi-1, glh-1, his-11, his-24, hmg-12, hpo-34, hrpk-1, hsp-1, hsp-25, hsp-3, icd-2, ifb-1, ifet-1, ifo-1, ketn-1, larp-1, let-75, let-805, lev-11, lfi-1, lin-22, lin-53, lmn-1, mec-7, mig-6, mlc-1, mlc-2, mlc-3, mlc-4, mlc-5, mup-2, myo-2, myo-3, myo-5, nkb-3, nmy-1, nmy-2, npp-13, nrfl-1, pab-1, pars-1, pat-10, pat-12, pqn-22, pqn-52, pqn-70, rpac-19, rpb-9, rpn-3, spc-1, sqd-1, tba-1, tbb-2, tcc-1, tln-1, tni-1, tni-3, tni-4, tnt-2, tsn-1, unc-15, unc-27, unc-54, unc-70, unc-87, vab-10, vgln-1, vig-1, vit-2, vit-5, vit-6, vrs-2 |

**Figure S5. PGN (*E. coli*)-binding proteins. Related to Figure 4.**
Bioinformatics analysis was performed based on a published dataset (Tian and Han, 2022) to annotate the subcellular localization of proteins in *C. elegans* that interact with *E. coli* OP50-derived PGN. A total of 168 PGN interacting proteins were classified into four categories: lysosomal, mitochondrial, ribosomal, and other/unclassified.

**Figure S6**

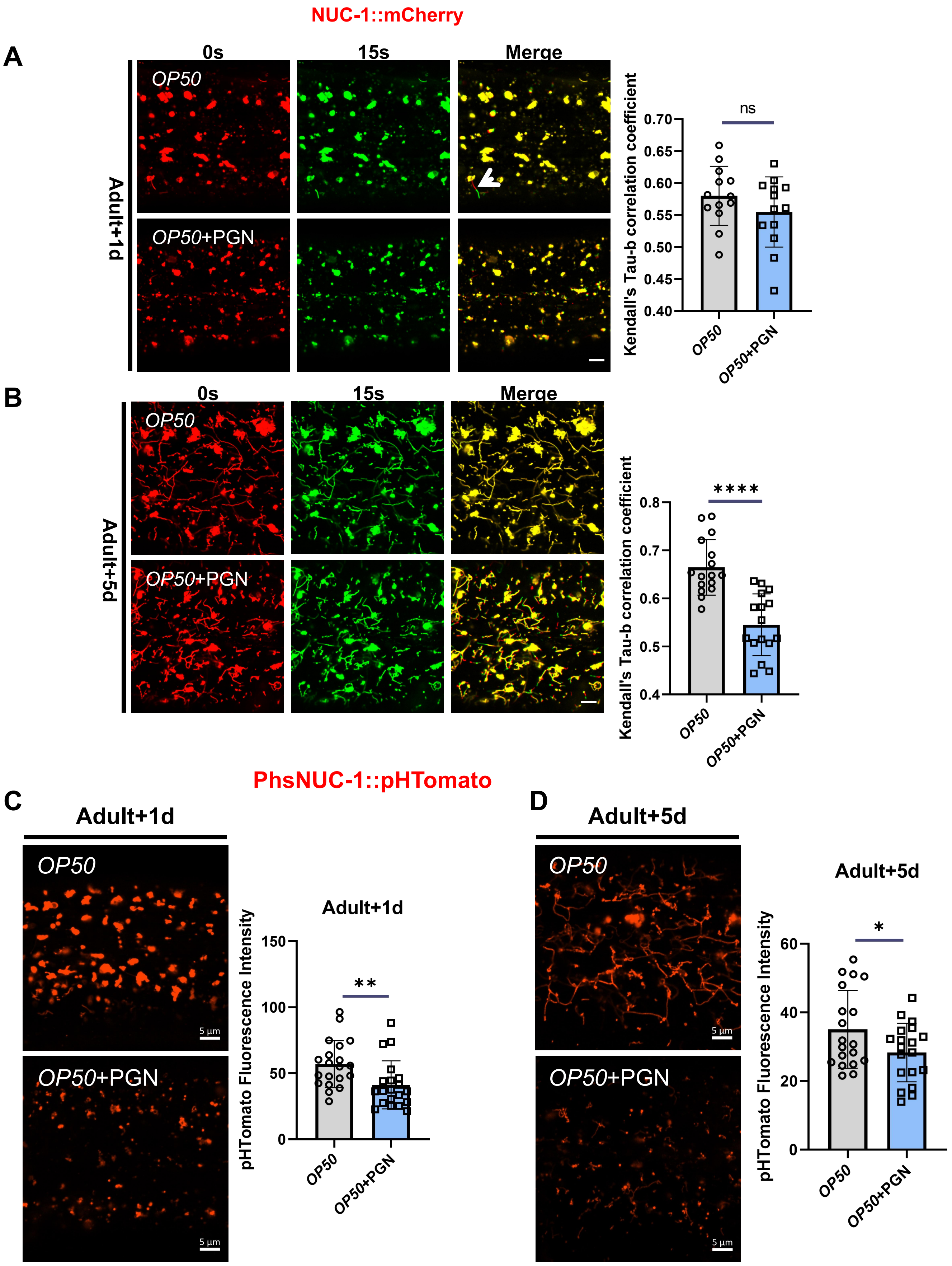


**Figure S6. PGN promotes lysosomal activity. Related to Figure 5.**

**(A, B)** Representative confocal images of NUC-1::mCHERRY lysosomes at 0 s and 15 s, and merged image, in Adult+1d(A) and Adult+5d(B) worms on OP50 or OP50+PGN. Bar graph shows Kendall's Tau-b correlation coefficients (lower = higher lysosomal dynamics). Scale bar, 5 μm. Data are mean ± SD. ns, not significant, ****P < 0.0001 (by unpaired two-tailed Student's t-tests). n ≥10 worms per group.

**(C, D)** Representative confocal images and quantification of pHTomato fluorescence intensity per lysosome in Adult+1d(C) and Adult+5d(D) worms on OP50 or OP50+PGN. Scale bar, 5 μm. Data are mean ± SD. *P < 0.05, **< 0.01 (by unpaired two-tailed Student's t-tests). n ≥ 20 worms per group.

#### Figure S7

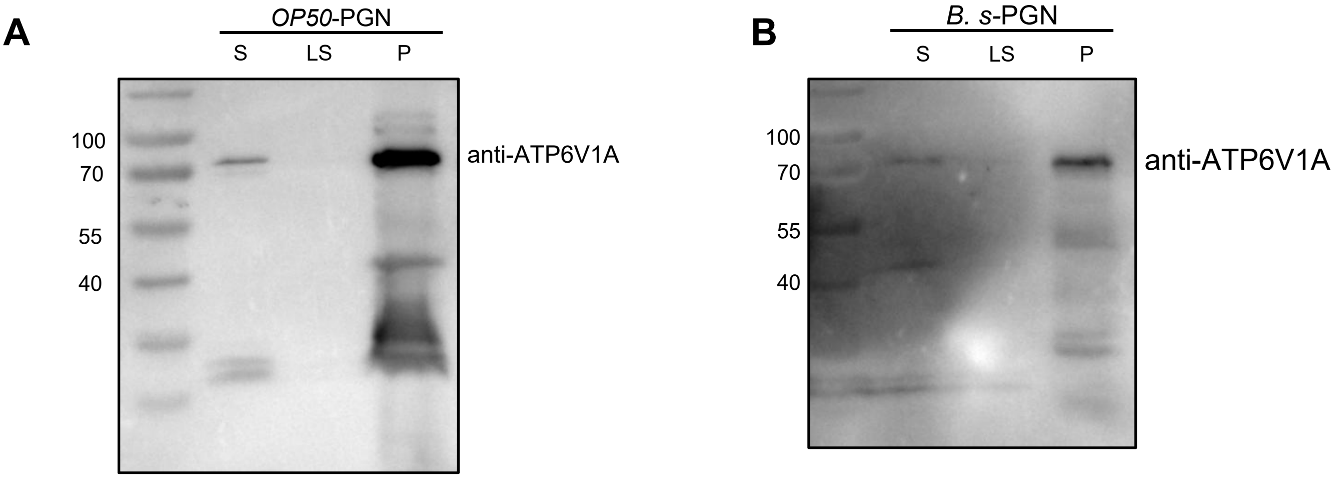


**Figure S7. PGN binds to ATP6V1A. Related to Figure 6.**

**(A, B)** Recombinant His-tagged ATP6V1A protein was expressed in E. coli and purified in vitro. The purified protein was incubated with *E. coli*-OP50-derived PGN (A) or *B. subtilis*-derived PGN (B) to verify their direct interaction in vitro.

All three fractions—post-incubation supernatant (S), final wash supernatant (LS), and washed pellet (P)—were analyzed by Western blotting (see more detail in methods section "PGN-binding protein assay"). The washed pellet (P) represents the PGN-bound protein fraction.
